# Groundfish with diverse life histories increase in size and abundance with proximity to spatial protections

**DOI:** 10.1101/2025.11.28.691246

**Authors:** Kyle L. Wilson, Alejandro Frid, Sean C. Anderson

## Abstract

Detecting positive responses to marine spatial protections, both *in situ* and as spillover effects, is critical for fisheries management yet difficult in practice. We leveraged 18 years of data on 28 groundfish species collected by four Indigenous Nations in Pacific Canada to assess ecological responses to spatially protected Rockfish Conservation Areas (RCAs). Body sizes and abundances were greatest inside RCAs and increased with proximity to RCAs at unprotected sites up to 160 km away. Abundance responses emerged faster (at younger RCA ages) for species with high r*_max_*, as predicted, but body size responses emerged faster for species with low r*_max_*, counter to predictions. Our study illustrates a general framework for jointly inferring *in situ* recovery and spillover benefits using a novel hierarchical geostatistical model that integrated long time series; observations made inside, near, and far from seven RCAs; and multiple species and survey methods spanning a 500-m depth range.

## 1. Introduction

Marine protected areas (MPAs) and other spatial fishery closures can increase the abundance and body sizes of exploited species, restore trophic structures, and enhance resilience (Baskett & Barnett 2015; Cheng *et al*. 2019; Kumagai *et al*. 2024; Micheli *et al*. 2012). The ecological consequences of restoring body sizes can be significant; within species, larger individuals have disproportionately higher fecundity per-capita (Marshall *et al*. 2021), and hold higher trophic positions than smaller individuals (Olson *et al*. 2020). Although these benefits are well-described inside MPAs compared to whole populations (Ovando *et al*. 2021), there is evidence that, as exploited populations recover within MPAs, adults or larvae “spillover” to unprotected areas to help rebuild fisheries (Barceló *et al*. 2021).

Interspecific variation in responses to MPAs can be significant (Claudet *et al*. 2008). For instance, longer-lived species, which have lower intrinsic population growth rates, r_max_, may require more years to respond than shorter-lived species (Jennings 2000). Additionally, adults of highly mobile species might cross MPA boundaries more frequently than sedentary ones, which may weaken their buildup of abundance within MPAs while potentially enhancing their contribution to spillover benefits at unprotected sites (Grüss *et al*. 2011).

The effectiveness of MPAs may also vary spatially according to their size, habitat suitability, connectivity to other MPAs, and enforcement of fisher compliance (Edgar *et al*. 2014). Spatial variation in fishing pressure—whether prior to MPA establishment and/or adjacent to the MPA after establishment—may amplify differences in population characteristics between protected and unprotected areas (Jaco & Steele 2020; Ziegler *et al*. 2022).

Rockfish (*Sebastes* spp.), a speciose genus of marine fishes found in the Northeast Pacific and primarily associated with structurally complex rocky reefs (Love *et al*. 2002), provide a compelling case study for examining concepts at the intersection of spatial protections and Indigenous governance. Indigenous Peoples living along the Northwest Coast of North America have been harvesting rockfishes sustainably for millennia (McKechnie & Moss 2016). Today, rockfish remain a traditional food for Indigenous Peoples, who strive to rebuild rockfish populations through MPAs (Reid *et al*. 2022) or other management measures (Petersen *et al*. 2020).

Rockfishes encompass diverse life histories and ecologies (Table 1). Some species have very long lifespans (e.g., ≥100 yrs) and therefore a lower r_max_ than shorter-lived species (Thorson 2020).

**Table 1.**
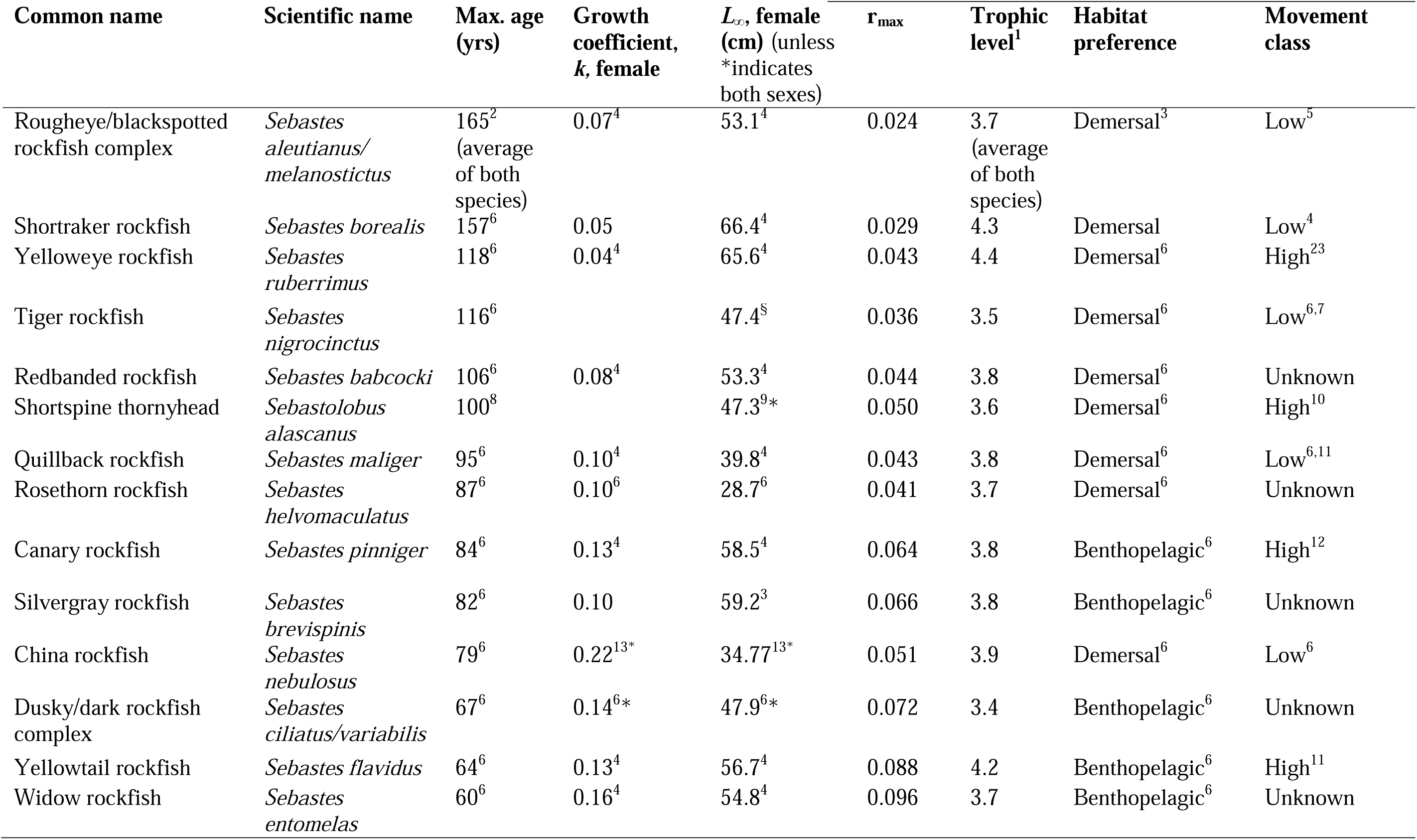
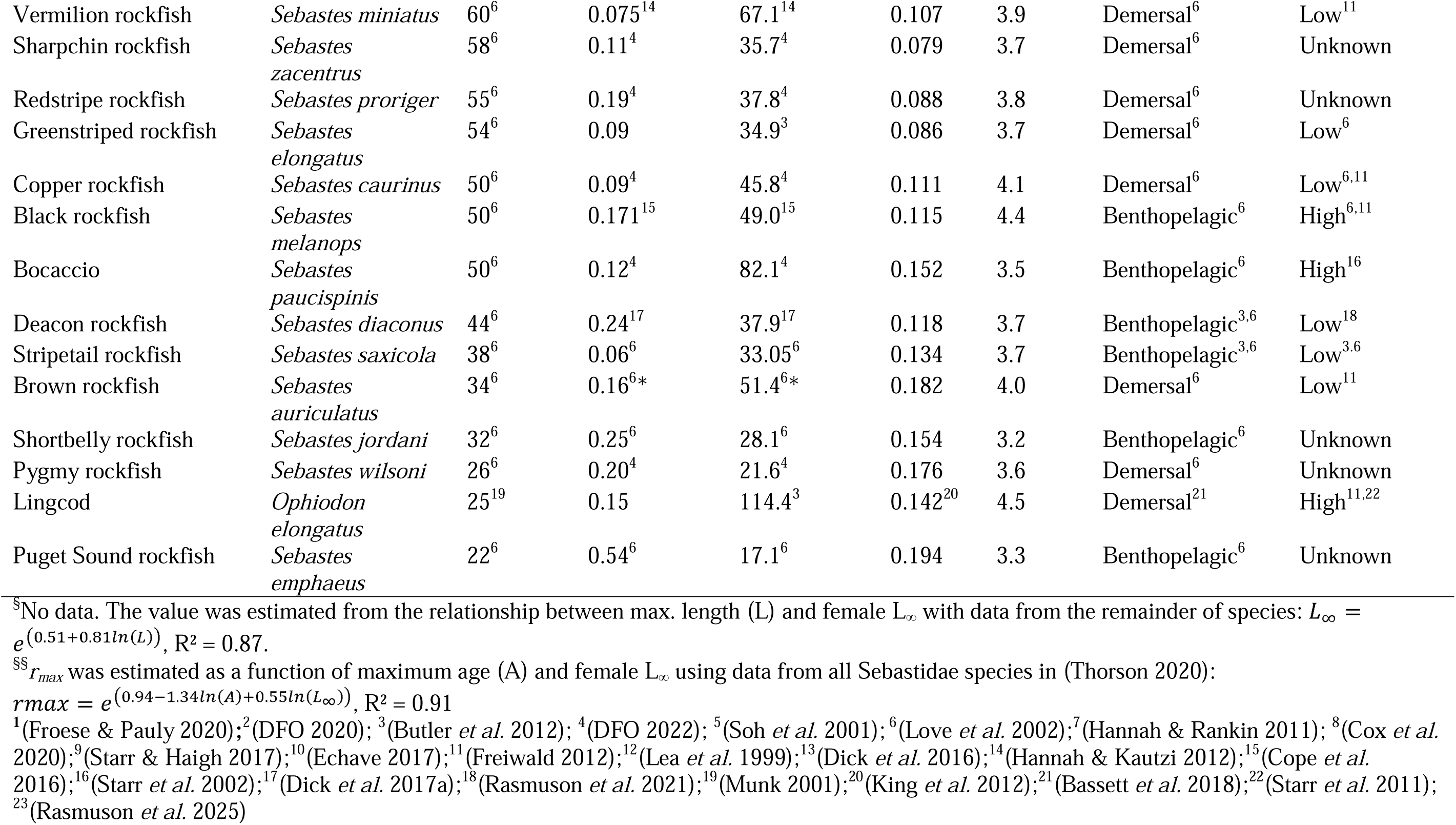
Species analyzed and their characteristics (ordered by maximum age). Superscripts indicate literature sources. Except for lingcod, intrinsic population growth rates, r_max_, were estimated from a generalized linear model^§§^ based on maximum age and *L* developed for all Sebastidae and based on global life history compendium sourced from (Thorson 2020). Maximum age for deacon rockfish are assumed to equal those for the closely related blue rockfish.

Rockfishes also encompass diverse ecological roles, from planktivores to predators at high trophic positions, and from sedentary demersal species to highly mobile benthopelagic species (Love *et al*. 2002). These traits may affect expectations for the extent to which MPAs restore different rockfish species (Jennings 2000; Moffitt *et al*. 2009), alter recovery timelines (Barceló *et al*. 2021; Kaplan *et al*. 2019), and affect food webs (Cheng *et al*. 2019; Olson *et al*. 2020).

In the 1980s, rockfishes in British Columbia (BC), Canada, underwent steep biomass and body size declines during a period of overexploitation (Eckert *et al*. 2018; Yamanaka & Logan 2010). By the early 2000s, changes to harvest management curtailed population declines and rebuilt biomass toward maximum sustainable yield (Anderson *et al*. 2021). From 2004–2007, to further aid recovery, Fisheries and Oceans Canada (DFO) implemented 162 Rockfish Conservation Areas (RCAs): long-term spatial closures that exclude bottom trawl, groundlines, and hook-and-line fisheries at locations cumulatively encompassing 20% of “good” rockfish habitats (Text S1). This trajectory of overexploitation followed by comprehensive management reforms, including spatial protections, parallels other Northeast Pacific groundfish fisheries and global trends (Hilborn *et al*. 2020; Keller *et al*. 2019; Warlick *et al*. 2018).

In 2023, four First Nations of BC’s Central Coast (Figure 1)—Wuikinuxv, Nuxalk, Kitasoo Xai’xais, and Haíłzaqv, who collaborate under the umbrella of the Central Coast Indigenous Resource Alliance (CCIRA)—completed a research program that amassed 13 years of fishery independent rockfish surveys collected over an 18-year period (Frid *et al*. 2016, 2018, 2021; McGreer *et al*. 2020). Working on behalf of CCIRA-member Nations, we analyzed this dataset, which spans seven RCAs and surrounding areas of BC’s Central Coast (Figure 1). Building on prior studies (Barceló *et al*. 2021; Jaco & Steele 2020; Kaplan *et al*. 2019), we tested (1) whether the abundance and body sizes of rockfish and co-occurring groundfish species increase with geographical proximity to RCAs (via spillover effects) and are greatest inside RCAs; and (2) whether these potential benefits of spatial protection increase over time at a faster rate (a) for species with faster life histories (higher r*_max_*) and (b) with the level of historical exploitation that preceded the establishment of RCAs. Additionally, we examined variation in the benefits of RCA proximity across all combinations of species and RCA locations.

**Figure 1.**
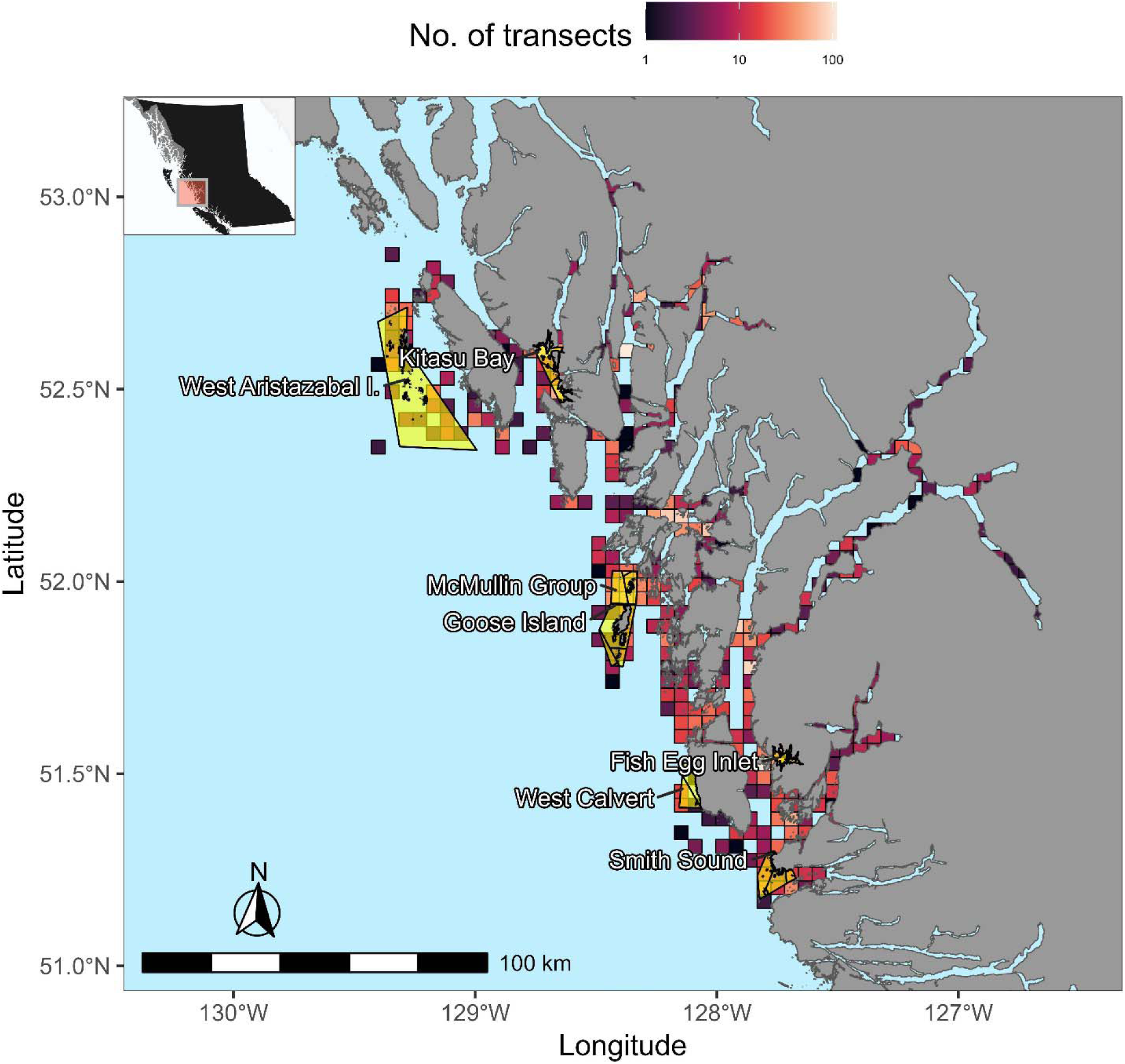
Map of Rockfish Conservation Areas (RCAs: yellow highlight) and transect locations in the Central Coast of British Columbia, Canada. Transect locations were aggregated within 16 km^2^ spatial units, which follows visualization protocols from the Central Coast First Nations (colours indicate the number of transect within spatial units). RCAs at Fish Egg Inlet, Goose Island, McMullin Group, and Smith Sound were established in 2004; the remainder of RCAs were established in 2005. The inset shows the study area location in Pacific Canada.

A key strength of our analysis is that it leveraged a long time series of observations made inside, near (<100 m), and very far (up to 160 km) from seven RCAs. We analyzed these observations with a directed acyclic graph (DAG)-informed structural causal inference framework (Arif & MacNeil 2023; Pearl 2009) and a novel geostatistical model (Anderson *et al*. 2025), integrating data from 28 species and multiple survey methods that spanned a 500-m depth range. Through this work, we illustrate a general framework for unifying inferences on *in situ* restoration and spillover benefits, providing a more holistic understanding of how different species respond to marine spatial protections.

## 2. Methods

### 2.1 Study design and data

CCIRA-member Nations hold Indigenous rights to their territories, where data were collected. Scientific staff who are members or employees of these Nations received approvals from rightsholders and were exempted from other permit requirements. DFO scientists partnered with First Nations to collect deep video survey data in their territories.

The data sets encompassed 26 rockfish species or species complexes, shortspine thornyhead (*Sebastolobus alascanus*), which belong to the same family as rockfishes (Sebastidae), and lingcod (*Ophiodon elongatus*) (Table 1). We focused on rockfishes and lingcod because they are explicit to the conservation objectives of RCAs (DFO 2026).

Data on relative abundances (count per unit effort, CPUE) were collected during 13 years of fieldwork (2006-2007, 2013-2023) using four fishery-independent methods (shallow diver transects, mid-depth video transects, deep video transects, and hook-and-line sampling) and one fishery-dependent method (sampling landings of Indigenous fisheries for food). As fish were counted only when present (caught or observed) during a survey, we applied depth and survey method selectivity criteria to infer true zeroes in the count data that may be biologically justified or reduce excessive zeroes that may be biologically unjustified. These methods are detailed in earlier publications (Frid *et al*. 2016, 2018, 2019, 2021; Gale *et al*. 2017; McGreer *et al*. 2020) and summarized in Table S1.

Additionally, body sizes (total length, in centimeters) were visually estimated for fish observed during dive transects (Text S2; Figure S1) or measured for specimens collected from hook-and-line sampling or Indigenous fisheries landings (Frid *et al*. 2016). The surveys included locations within 7 RCAs established in 2004 or 2005 (Figure 1).

Most rockfishes and lingcod associate with structurally complex rocky reefs (Love *et al*. 2002). Surveys, therefore, targeted rocky reefs and did not randomly sample all available bottom types, which include soft substrates. For dive surveys, surveyed reefs were limited to depths of ≤35 m (SCUBA limits) while other methods sampled deeper depths (Table S1). Within these criteria, sites were chosen using Indigenous knowledge, local knowledge, bathymetric models (Haggarty & Yamanaka 2018), and (to a lesser extent) ad hoc exploration. Rocky reefs vary in structural complexity and are fragmented by soft-bottom habitats; analyses controlled for variability in habitat structural complexity.

Each transect or sampling event had a spatial resolution of ≤120 m^2^ (480 m^3^ for dive transects: Table S1). For each location, we collected or compiled data on potential drivers and covariates that included habitat structural complexity, bottom depth, time of year (measured as 26 two-week bins), sampling gear, sampling effort, and the name of and distance (km by water avoiding land) to the nearest RCA, with a distance of 0 corresponding to locations inside RCAs. Overall, 2,493 transects (of 4,307 total) observed ≥1 species of groundfish (1,178 dive transects; 1,017 mid-depth video transects; 126 deep video transects; and 172 hook-and-line transects) and a total of 139,402 groundfish were enumerated. In addition, 1,353 transects measured total lengths of groundfish (1,175 from dive transects and 178 from hook-and-line transects). We recorded a total of 15,906 unique size cohorts, i.e., the number of fish within a school of a particular species (≥1 fish of that species) observed at a particular total length.

DFO provided commercial fishery catch data covering a 1-km radius around each transect location, spanning from 1996 to 2005. For each species and location, we summed the total catch prior to RCA establishment (either 2004 or 2005) as a measure of cumulative harvest at that location for the pre-RCA period. We assumed zero catch for locations with missing data. Records of recreational fishing pressure (legal or illegal) were unavailable, and our analyses may underestimate total cumulative harvest at some locations.

### 2.2 Analytical approach

We developed a directed acyclic graph, or DAG (Arif & MacNeil 2023; Byrnes & Dee 2025; Pearl 2009), representing hypothetical causal relationships of RCA proximity and RCA age to CPUE and total length. Like other causal diagrams, the DAG encodes assumptions about plausible pathways, which we make transparent in Figure S2 and Text S3. The DAG’s structure was reflected in the design of our geostatistical generalized linear mixed-effects models (GLMMs; Table S2) fitted with the *sdmTMB* package in R version 4.6.1 (Anderson *et al*. 2025; R Core Team 2026). The spatial GLMMs examined the role of the primary drivers of interest: proximity to RCA, RCA age, cumulative harvest up to the point of RCA establishment (pre-RCA harvest, for brevity), and species life histories (i.e., r*_max_*). The DAG implies an adjustment set comprising variables that influence both RCA placement and responses: pre-RCA harvest, because RCAs were established in areas with particular fishing histories; depth and habitat complexity, which shape rockfish distributions and informed RCA boundaries; and survey gear and time of year, which affect observation rather than the underlying process but are confounded with location and season of sampling (Carlson & Barr 1977; Love *et al*. 2002). Given the proposed DAG, conditioning on these variables would block the non-causal pathways between RCA proximity and the responses, leaving the association attributable to spatial protection. Latent spatial variation was accounted for by random fields (see Tables S5, S6).

Cumulative harvest enters the DAG differently depending on when it occurred: harvest before RCA establishment is a common cause of both RCA placement and the responses, whereas harvest after establishment lies on the causal pathway from protection to the responses. Although cumulative harvest after RCA establishment may affect responses to spatial protection (Ziegler *et al*. 2022), we do not include it as a covariate because, under our assumed DAG, it functions as a post-treatment mediator and conditioning on it would block part of the causal effect of interest (Byrnes & Dee 2025; Cinelli *et al*. 2024). Additionally, species mobility may affect responses to spatial protection (Moffitt *et al*. 2009), but movement information was unavailable for 10 of 28 species (Table 1). Accordingly, GLMMs excluded harvest after RCA establishment and species mobility.

We expected CPUE and total length to be greater, on average, inside RCAs and to decline with increasing distances from RCAs. Hence, we modelled spatial protection as the distance of each location to the nearest RCA, where 0 km delineated locations inside RCAs. We recorded distances outside RCAs as negative values such that a positive influence of RCA “proximity” on the response variable would correspond to a positive effect. We also expected spatial protection benefits to manifest faster for species with higher r*_max_* and with higher pre-RCA harvests. Hence, we modelled pairwise interactions between: (1) r_max_ and RCA age, (2) r_max_ and RCA proximity, (3) RCA proximity and pre-RCA harvest, and (4) RCA distance and RCA age.

### 2.3 Geostatistical model

We modelled the observed counts *C* of the *i*th sample in transect *j* of species *k* at location *s* sampled by gear *g* in year *t* using a negative binomial distribution (NB2: Hilbe 2011) such that:

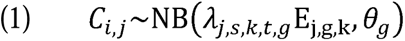

where *θ_g_* represented the dispersion parameter for survey gear *g* and the expected count was modelled as the product of the expected catch-per-unit effort *λ_i,s,k,t,p_* and an offset for transect effort E_j,g,k_ (effort units for dive and towed video: m^2^; hook-and-line or long-line: total hook-hours). For video transects, we bias-corrected effort by species’ chase probability (P_k_) to account for potential attraction to the camera’s parallel laser beams, which would inflate capture efficiency for those species, such that 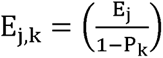 (Frid *et al*. 2020). We described the expected CPUE *λ* using a linear predictor with a log link such that:

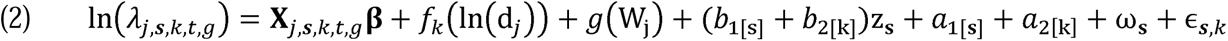

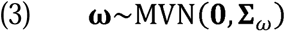

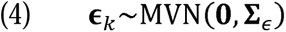

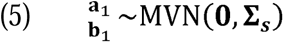

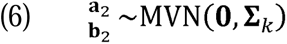

where **X***_J,s,k,t,g_* represented a model matrix of predictors that were multiplied by a vector of fixed-effect coefficients **β**, *f*_k_(ln (d*_J_*)) represented a penalized smoother for the natural log of depth (m) that varied for groups of species that occupy similar depth ranges (Figure S3), *g*(W_j_) represented a penalized smoother for the bi-weekly time of year, *b*_2[k]_ and *b*_1[s]_ represented a random slope of the proximity to the nearest RCA (denoted by z_s_) stratified by species *k* and the RCA nearest to location *s*, respectively, *a*_1[s]_ and *a*_2[k]_ represented the random intercept for species *k* and the nearest RCA to location *s*, respectively, and ω_s_ and ɛ*_s,k_* represented overall spatial and species-specific spatial random effects, respectively. Random intercepts and slopes for species and RCAs were modeled jointly as bivariate normal distributions (Eqs. 5, 6), where 2×2 covariance matrices **Σ***_k_* and **Σ***_s_* capture the variance of each component and their correlation for species and RCAs, respectively. Spatial and species-specific spatial random effects were assumed to be drawn from Gaussian Markov random fields with covariance matrices (inverse precision matrices) **Σ***_ω_* and **Σ***_∈_*, each structured to approximate a Gaussian random field with a Matérn covariance function, to account for spatially correlated residual variation (Lindgren *et al*. 2011). We modelled all spatial random effects using a physical barrier (Bakka *et al*. 2019) that accounts for dispersal and movement barriers among the islands and fjords typical of the study area. Our barrier mesh implementation assumed that spatial correlation decayed 10 times faster over land than water.

Similarly, we modelled the observed total length *L* of the *i*th size cohort sampled in transect *j* of species *k* at location *s* sampled by gear *g* in year *t* using a gamma distribution (specified as shape, scale) with mean µ (with a log link) and variance µ^2^ /*ϕ* such that:

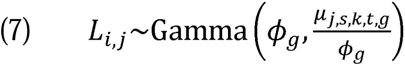

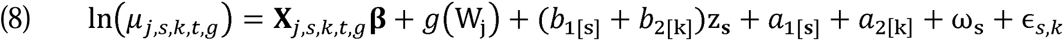

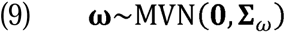

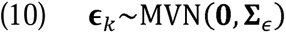

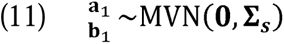

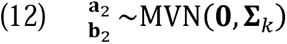

with parameter descriptions similar to the count model. For total length, we modelled a species-specific linear effect of the natural log of depth (Figure S4) rather than a penalized smoother because of issues with model convergence. During dive surveys, divers recorded benthopelagic species, which may be encountered in large schools, as the number of fish sharing the same estimated total length, so a single transect could yield several size cohorts. We weighted each observation’s log-likelihood by its share of that species’ count in that transect, normalised within transect and species so that the mean weight was 1 and each transect’s weights summed to its number of size cohorts (see ‘data weighting’ examples in Table S3 and S4). A cohort’s influence therefore depends on its share of the transect count rather than on absolute abundance, and large schools do not dominate the total length model. These analyses excluded fish with lengths <20% of *L*_∞_, which are unlikely to have recruited to the adult population (Lorenzen & Camp 2019), and species with <100 observations of total length.

### 2.4 Model validation & inference

Each GLMM was fit by maximizing the marginal log-likelihood using TMB, which integrates over random effects via the Laplace approximation (Kristensen *et al*. 2016). We used several complementary methods to diagnose model suitability. First, we ensured that each model passed a suite of checks and criteria that indicated consistency with model convergence, e.g., positive definite Hessian matrices, all fixed effect absolute log likelihood gradients < 0.001, and estimated standard deviations > 0.01 (i.e., not on a boundary) (Anderson *et al*. 2025). We compared observed versus fitted values for every species-transect record of both CPUE and total length as a goodness-of-fit diagnostic (Figure S6). We also examined simulation-based randomized quantile residuals using the package *DHARMa* (Hartig 2024) (Figures S7-S10). Specifically, we took a single draw from the implied random effect MVN distribution with fixed effects at their maximum likelihood estimates (Waagepetersen 2006) and applied 1,000 iterations of observation error. We then compared the distribution of these predictions to the observations for each survey gear and species. For each response variable (CPUE or body size), we characterised the direction and strength of RCA contributions to spatial protection for each species-RCA combination using effect sizes (i.e., multiplicative change in the response variable per 5-km increase in RCA proximity), 95% Confidence Intervals (CIs), and Pr(β>0) — the probability of a positive effect — which indicates the strength of evidence for a directional response on a continuous scale. Both 95% CIs and Pr(β>0) were constructed from 10,000 simulations of the fixed and random effects from the model’s joint precision matrix. We report Pr(β>0) alongside CIs for each species-RCA combination to distinguish the consistency of an effect’s direction from the precision of its magnitude, and we interpret these quantities directly rather than applying a threshold rule. Marginal effects were computed using the *emmeans* package (Lenth 2025). Lastly, we compute partial working residuals on the log-link scale for each observation of CPUE and total length to allow for graphical assessment of the marginal effect of RCA proximity to the empirical data.

We conducted a suite of sensitivity tests to determine whether estimates of the main effect of RCA proximity were robust against alternative model structures. For both responses, we tested whether the dispersion parameter was stratified by survey gear, barrier versus non-barrier mesh, and the cutoff resolution of the mesh. For CPUE, we additionally tested the NB1 (linear variance) versus NB2 (quadratic) parameterization of the negative binomial distribution (Hilbe 2011) and the treatment of extreme schooling observations (>1,000 schooling individuals). For total length, we tested a Gamma versus lognormal distribution and four data-weighting approaches: (a) the base model, (b) equal weights per size cohort, (c) weights proportional to expected CPUE, normalised within species, and (d) equal weights per school (Figure S5).

## 3. Results

Overall, the marginal effect of RCA proximity on total length was weak but consistently positive (0.05; 95% CI: 0.02 – 0.07; Pr(β>0) = 1.00), whereas the effect on CPUE was larger but more uncertain (0.58; 95% CI: -0.04 – 1.20; Pr(β>0) = 0.97; see Tables S5 and S6 for estimated coefficients and variance parameters). These effects indicate that, for every 5-km increase in RCA proximity, the average species increased in total length and CPUE by factors of 1.01 (95% CI: 1.00 – 1.02) and 1.11 (95% CI: 0.99 – 1.25), respectively (Figures 2a, 3a). Variation around these average responses depended on differences among species-level r_max_, RCA ages, and pre-RCA harvest (Figures 2b, 3b). Estimates of RCA proximity effects on CPUE and total length were consistent across most alternative model assumptions (Figure S5). However, the effect of RCA proximity on CPUE was sensitive to the resolution of the barrier mesh, with point estimates ranging from 0.36 to 0.79, corresponding to CPUE increases of 7–15% per 5-km increase in RCA proximity.

**Figure 2.**
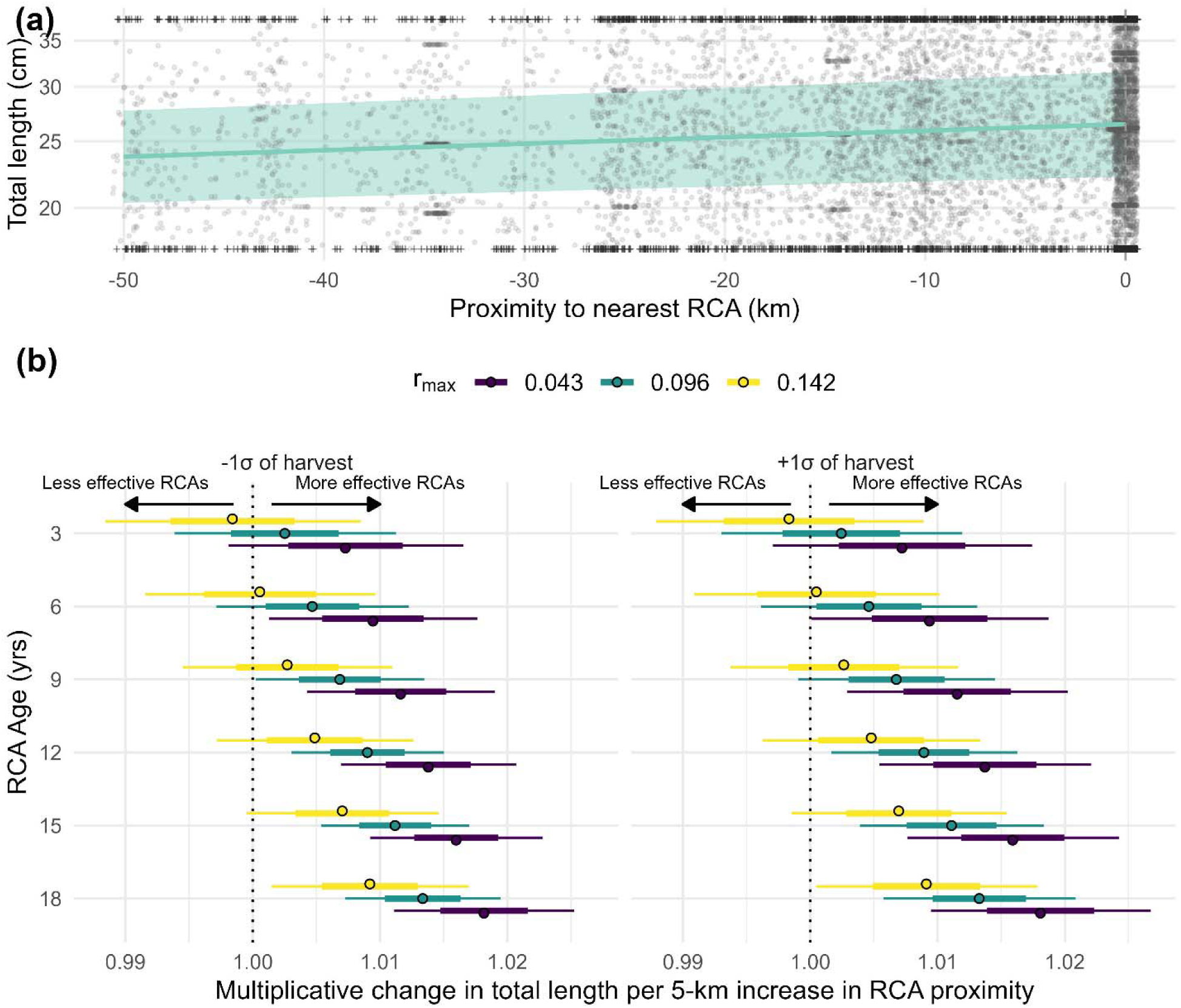
(a) Marginal effect of RCA proximity on total length (i.e., with remaining covariates at their average values); shading depicts 95% confidence interval. Grey points indicate partial working residuals; crosses indicate working residuals that were outside the upper or lower limit of the plotted range. The dashed line marks the RCA boundary, where observations were made inside protected areas. (b) Conditional effect sizes of RCA proximity (per 5-km increase in RCA proximity) on total length in response to interactions with RCA age, r*_max_*, and pre-RCA cumulative harvest. Low (left panel) and high harvests (right panel) are standardized as ±1SD of species-specific harvests (e.g., 0kg km^-2^ and 358 kg km^-2^ for quillback rockfish). Thinner outer bars and thicker inner bars delineate, respectively, 95% and 66% confidence intervals. Categories of r_max_ are based on the following species: yelloweye and quillback rockfish for low (r_max_= 0.043); widow rockfish for mid (r_max_= 0.096); lingcod for high (r_max_=0.142). The range of RCA ages considered matches the empirical data.

The positive effect of RCA proximity on total length strengthened as RCAs became older (Figure 2b; RCA age × RCA proximity: 0.01; 95% CI: 0.00 – 0.02; Pr(β>0) >0.99). This effect manifested earlier for species with lower r*_max_* (r*_max_* × RCA proximity: -0.02; 95% CI: -0.04 – 0.00; Pr(β>0) = 0.02), while pre-RCA harvest had little influence (pre-RCA harvest × RCA proximity: 0.00; 95% CI: -0.01 – 0.01; Pr(β>0) = 0.48).

The positive effects of RCA proximity on CPUE also strengthened as RCAs became older but were somewhat uncertain (Figure 3b, RCA age × RCA proximity: 0.06; 95% CI: -0.01 – 0.14; Pr(β>0) = 0.95). Positive effects manifested earlier for species with higher r*_max_* — the opposite of body size responses (r*_max_*× RCA proximity: 0.28; 95% CI: -0.14 – 0.70; Pr(β>0) = 0.91) — and weakened with higher pre-RCA cumulative harvests (pre-RCA harvest × RCA proximity: -0.073; 95% CI: -0.188 – 0.042; Pr(β>0) = 0.11). At lower pre-RCA harvests, effect sizes for species with mid r*_max_* increased steadily with RCA age, while estimates for species with low r*_max_* remained positive but poorly resolved even after 18 years (Figure 3b).

**Figure 3.**
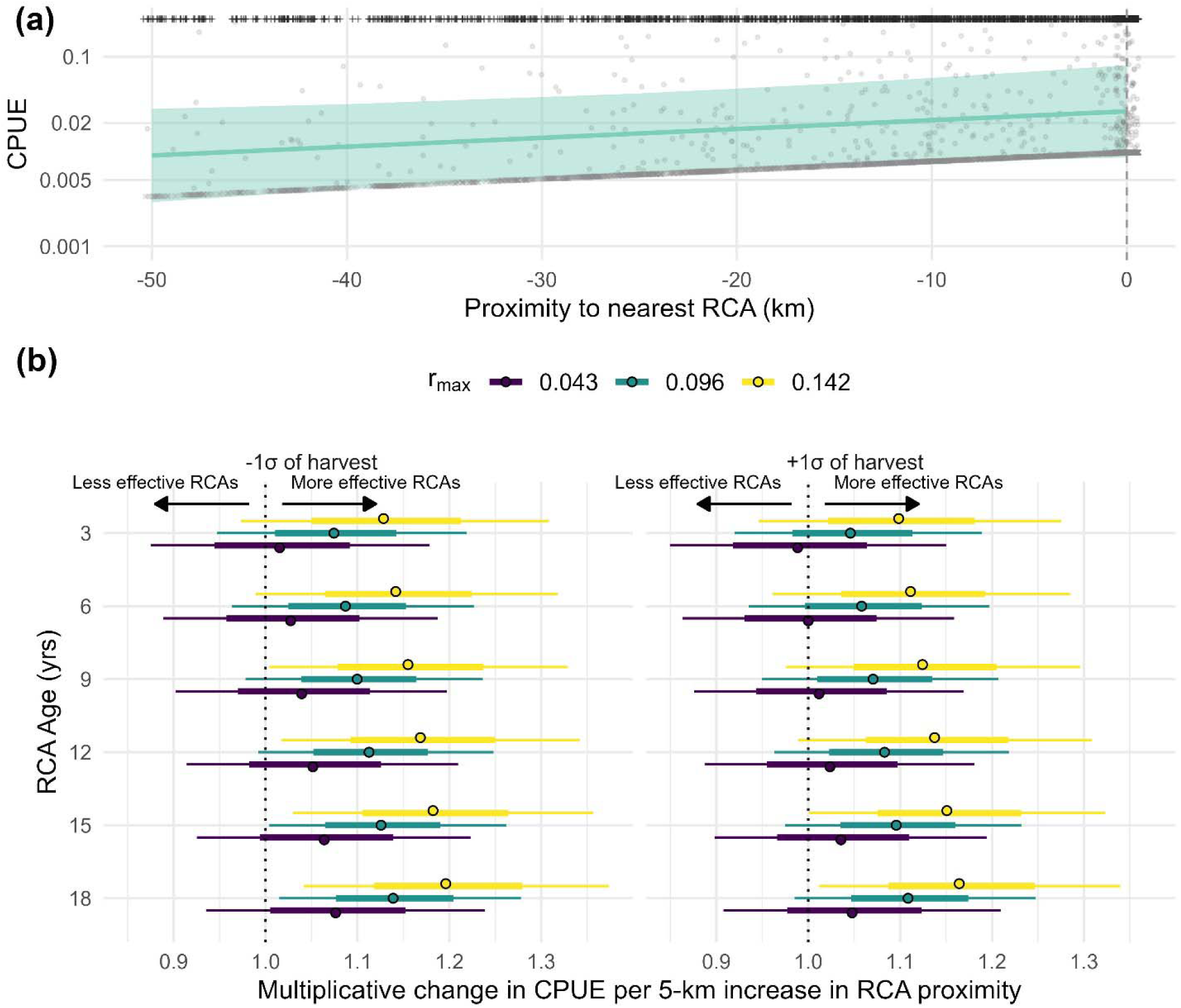
a) Marginal effect of RCA proximity on CPUE (i.e., with remaining covariates at their average values); shading depicts the 95% confidence interval. Grey points indicate partial working residuals; crosses indicate residuals that were outside the upper limit of the plotted range. Partial residuals for all zero counts fall along a parallel band beneath the ribbon. The dashed line marks the RCA boundary, where observations were made inside protected areas. (b) Conditional effect sizes of RCA proximity (per 5-km increase in RCA proximity) on CPUE in response to interactions with RCA age, r*_max_*, and pre-RCA cumulative harvest. See caption to Figure 2 for details.

The effects of RCA proximity varied between RCA locations and species (Figure 4). The point estimates of RCA proximity were positive for 100% of 98 species-RCA combinations for total length (median Pr(β>0)=0.99) and 69% of 179 species-RCA combinations for CPUE (median Pr(β>0)=0.78). There was strong support for positive effects (Pr(β>0) ≥ 0.90) for 89% of species-RCA combinations for total length and 35% of combinations for CPUE; conversely, there was strong support for negative effects (Pr(β>0) ≤ 0.10) for 0% and 7% of combinations, respectively. Effect sizes were highest for lingcod (total length; a factor of 1.02 per 5-km increase in RCA proximity; 95% CI: 1.01 – 1.03) and vermilion rockfish (CPUE; a factor of 1.59; 95% CI: 1.29 – 1.96).

**Figure 4.**
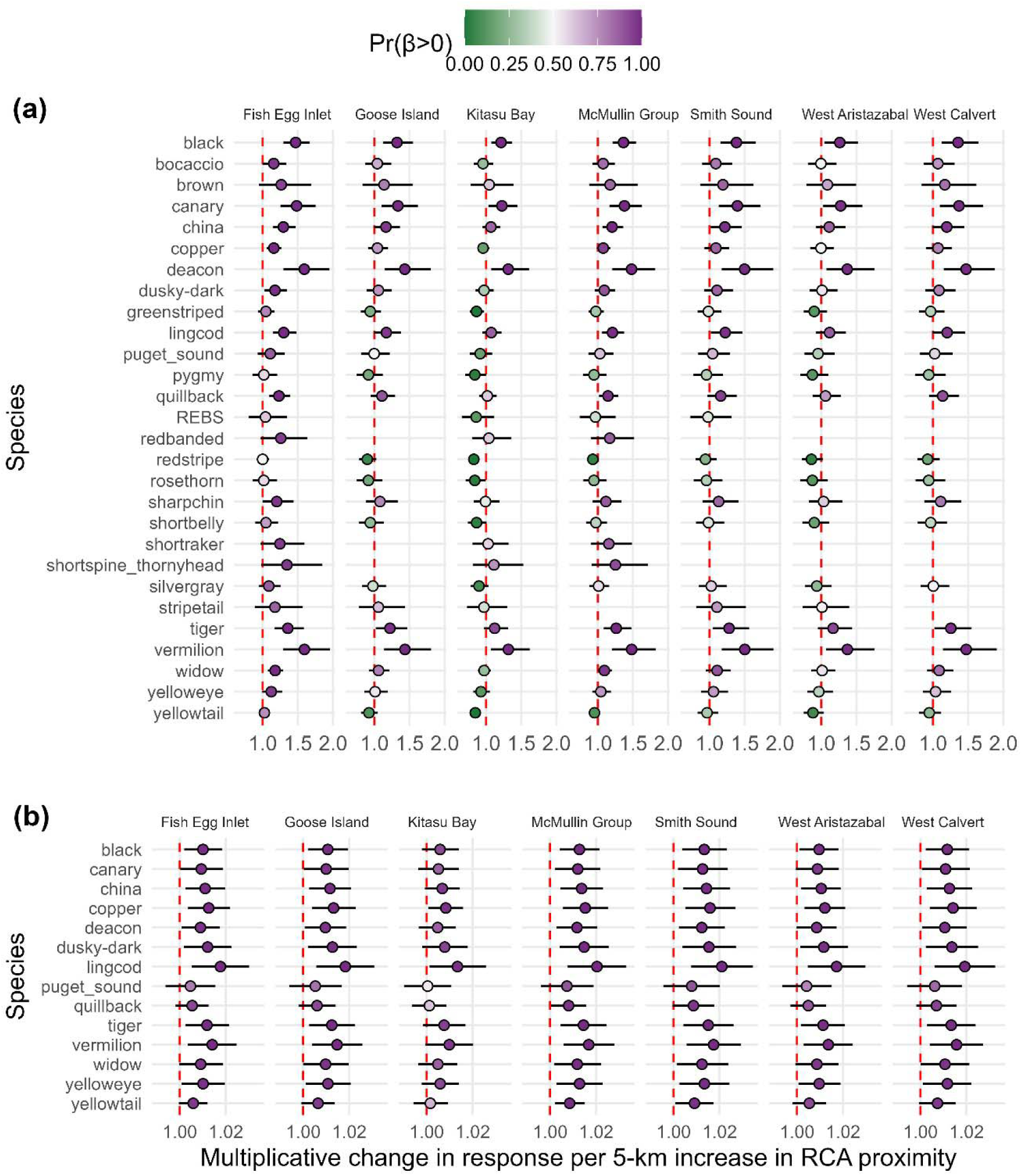
Mean and 95% confidence intervals for the estimated effect sizes of RCA proximity on (a) relative abundance (i.e., CPUE) and (b) total length, by RCA location (upper X-axes labels) and species (Y-axes labels). Effect sizes were calculated by summing the estimated fixed and random effects in log space and transforming to the response scale, per 5-km increase in RCA proximity; uncertainty intervals were constructed by simulating 1,000 draws from the estimated joint precision matrix of the spatial GLMM. Point colours indicate Pr(β>0), the proportion of draws in which the effect was positive. The vertical dashed line indicates the zero line for visual aid. The term REBS represents the rougheye/blackspotted rockfish complex.

Some temporal patterns uncovered by our data (i.e., effects of RCA age relative to implementation) may reflect drivers that were partially independent of spatial protection, and which elicited different responses across species (Figure 5). Over time, CPUE increased for species with low r_max_ inside and outside RCAs, though faster inside; decreased for species with high r_max_ both inside and outside, though more slowly inside; and diverged only for species with mid r_max_, increasing inside while declining outside (Figure 5a). Total length decreased over time for species with low and mid r_max_ both inside and outside RCAs, though more slowly inside and faster overall for low r_max_. With high r*_max_*, total length increased inside RCAs with changed little outside (Figure 5b).

**Figure 5.**
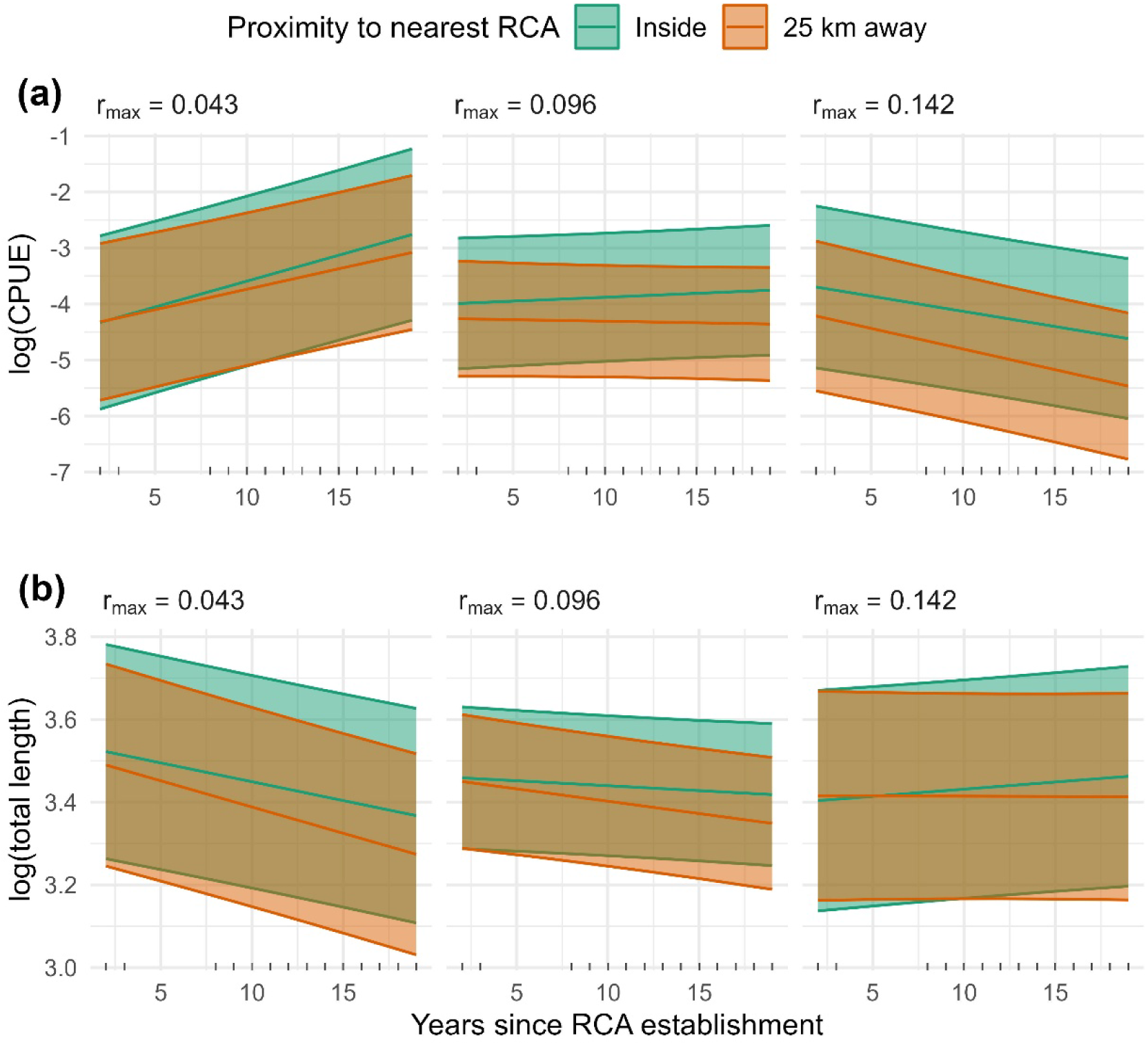
Marginal effects of RCA proximity on (a) relative abundance, log(CPUE), and body size, log(total length), by selected categories of RCA proximity and r*_max_*. Shaded ranges indicate 95% confidence intervals. Tick marks along the lower axis show the RCA age at each surveyed transect.

Total length and CPUE also responded to depth (Figures S3, S4), habitat complexity, survey gear, and time of year (Table S5). The estimated Matérn ranges (the distance spatial correlation becomes negligible) were 10 km and 17 km for total length and CPUE, respectively. We estimated large residual spatial variation for total length and CPUE within and across species (Table S6). Estimated spatial random fields represented a large portion of model complexity as measured by effective degrees of freedom (Table S7).

## Discussion

A large body of work provides evidence that marine spatial protections can restore larger body sizes and greater abundances within protected areas (Baskett & Barnett 2015; Bosch *et al*. 2022). These studies often have a paired sampling design that compares population characteristics inside MPAs and at nearby unprotected areas (Micheli *et al*. 2012; White *et al*. 2021; Willis *et al*. 2003). A second body of work provides evidence that larvae or adults that originate within MPAs move into fished areas where they boost abundance and potentially improve fisheries catches, often far from MPAs (Barceló *et al*. 2021; Franceschini *et al*. 2024; Le Port *et al*. 2017). Our results, which derive from observations made inside RCAs and across a large range of distances from them (Figures S11, S15), simultaneously address *in situ* restoration and spillover benefits. Furthermore, adult spillover is often examined as shifts in catches or abundance at different distances from spatially protected areas (Halpern *et al*. 2009; Murawski *et al*. 2005), yet our results include a less common documentation of larger body sizes extending into fished areas (Franceschini *et al*. 2024).

Consistent with our predictions, RCA proximity had positive effects on the body size and relative abundance of rockfishes and related species. Although the proximity effect was larger in magnitude (but also more uncertain) for relative abundance than body size, the role of RCA proximity in rebuilding or maintaining size-structures is particularly important because the hyperallometry of size-fecundity relationship is strong for rockfish (Dick *et al*. 2017b). That is, subtle increases in body sizes in response to spatial protections may manifest as disproportionate increases in larval production. Marshall *et al*. (2019) suggests that hyperallometric fecundity can underestimate the reproductive contribution of fish inside MPAs due to Jensen’s inequality. For instance, average total length increased by 1.1% per 5 km-increase in RCA proximity. Applying a hyperallometric size-fecundity exponent of 1.18 (Barneche *et al*. 2018) and a Jensen’s inequality correction (Marshall *et al*. 2019), this implies an increase in per-capita fecundity of ≈1.6% moving from 5-km outside to inside an RCA, and ≈14% across a 50-km gradient. Among other benefits, increased annual larval production could enhance resilience to environmental perturbations (Micheli *et al*. 2012). Estimates of the effect of RCA proximity on body size were consistent among species, though some sites far from RCAs are in fjords where environmental constraints on maximum body size may be greater (West *et al*. 2014). Still, these results could help to build spatially explicit population models estimating recovery timelines across a wide range of distances from RCAs (Ovando *et al*. 2021).

We also predicted that the positive effects of RCA proximity would require less time to manifest for species with faster life histories (i.e., higher r_max_), but this prediction was supported only for abundance responses. Contrary to predictions, the positive effects of RCA proximity on body size manifested at earlier RCA ages for species with lower r_max_. The tension between these two sets of results is interesting. One potential explanation relates to species mobilities, which are unaccounted for in our model due to missing information for 10 of 28 species. Of the species with mobility information, those with higher r_max_ tend to be more mobile (Table 1) and, therefore, more likely to leave protected areas before reaching larger body sizes (Moffitt *et al*. 2009); these characteristics increase their risk of size-truncation from fisheries exploitation (Yan *et al*. 2026). Additionally, responses to ocean warming – which can curtail maximum body sizes (Cheung *et al*. 2012) – could potentially vary between species that differ in r_max_. More generally, r_max_ cannot encapsulate all species differences in traits expected to affect responses to spatial protections. For instance, the von Bertalanffy growth coefficient, *k*, also affects these responses (Kaplan *et al*. 2019), yet variation in r_max_ explains only 35% of the variation in *k* for species in our study (Figure S12).

The effects of r_max_ on responses to RCA age must be read against broader temporal patterns that may reflect drivers other than spatial protection. Inside and outside RCAs, the abundance of species with high r_max_ and the body sizes of species with low and mid r_max_ *declined* over time while the abundance of species with low r_max_ *increased* over time. Hypotheses for these patterns include species differences in (1) responses to the combined effects of environmental change and the legacies (genetic or phenotypic) of overfishing (Heino *et al*. 2015; Yan *et al*. 2026); (2) vulnerability to illegal fishing in RCAs and legal recreational fishing in unprotected areas; and (3) correlates between r_max_ and mobility discussed above. Regardless of the mechanisms, we found that body size and abundance increased faster or declined more slowly inside than outside RCAs, suggesting that spatial protection may have lessened impacts of environmental change (Ziegler *et al*. 2023).

Counter to expectation (Jaco & Steele 2020), increased fishery harvests prior to RCA establishment weakened or had a negligible effect on, respectively, abundance and body size responses to spatial protection over time. A potential explanation is that sites adjacent to RCAs with high pre-RCA harvests continued to experience higher harvests after RCA establishment (Figures S13, S14). These sites may require more time to accrue spillover benefits associated with RCA proximity than sites with lower harvests either pre- or post-RCA establishment. Alternatively, sites with higher pre-RCA harvest may have been more productive and less depleted: pre-RCA harvest had a consistent, positive main effect on CPUE and total length (Table S5), suggesting that historical fishing concentrated where fish were larger and more abundant.

Responses to RCA proximity were unexpected in some cases. Strong responses, whether positive or negative, were absent for most CPUE combinations (58%) but few for total length (11%). This distribution of RCA proximity effects cautions that some effects, whether positive or negative, could potentially represent statistical noise. However, strong positives outnumbered strong negatives by 5:1 for CPUE results, and there were no strong negatives for total length responses. Thus, positives were unlikely to represent the upper half of a noise distribution (Figure 4). Potential ecological explanations for a null response to RCA proximity include little to no harvest during the pre- and/or post RCA periods (Figures S13, S14), which implies an expected effect near zero rather than a decline. For example, nine of the 11 species lacking strong positives for abundance responses to all RCAs experienced relatively low harvests during both pre- and post-RCA implementation periods (Table S8). The recovery of larger-bodied species inside RCAs may increase predation upon smaller-bodied species (Beaudreau & Essington 2011), which may underlie the weak or strongly negative abundance responses of some smaller-bodied species (Table S8). Effort displaced from RCAs may also concentrate outside their boundaries (Halpern *et al*. 2009), which would depress abundance at nearby unprotected sites and generate a negative association with RCA proximity; this mechanism may be more plausible for targeted, schooling species such as yellowtail and redstripe rockfish than for the small-bodied species described above.

Our inferences depended on data collected after RCAs were established. Without a BACI design (sensu Underwood 1991), we made untestable causal assumptions (Correia *et al*. 2026): that environmental controls in our statistical model (depth, habitat complexity, and spatial random effects) captured all pre-existing habitat gradients prior to RCA establishment, and that post-establishment harvest did not reflect an exogenous shock (e.g., policy or market change) that independently affected rockfish ecologies (Ferraro *et al*. 2019). These limitations highlight the need to scrutinize alternative explanations of our findings. For example, an environmental decline might concentrate fish near high-quality habitat that coincide with RCAs, which could confound an environmental effect with one of spatial protection. This scenario, however, seems unlikely for our study, as the RCA network was designed to protect only 20% of “good” rockfish habitats, leaving 80% of that habitat unprotected. In another scenario, RCAs are effective but adult movement and larval dispersal are high enough to dampen the spillover signal by rebuilding abundance inside and outside RCAs to similar levels, which may underestimate RCA effects if we had studied only sites within and adjacent to RCAs. However, our analyses examined the effects of RCA proximity for unprotected sites spanning a 160-km-range of distances from the nearest RCA boundary. Had a BACI design been possible, the RCA effects that we detected might have been stronger.

Regardless of spatial and interspecific variability, our results provide consistent but variable evidence that body sizes and abundances were, on average, greatest inside RCAs and increased with proximity to RCAs at unprotected sites. These benefits increased over time. Understanding the timescales of these effects, and the ability to measure them, is essential for management. If our study had been shorter than the 18-year post-RCA period, the value of RCAs and their validation as management tools would have been difficult to resolve from empirical data (Haggarty *et al*. 2016). Our results, therefore, serve as strong encouragement for management agencies to commit resources for robust, stable, and *long-term* monitoring programs designed to examine the effectiveness of spatial protections (see DFO 2026).

More generally, distinguishing signal from noise here depended on the length of the time series, the inclusion of multiple protected areas spanning a wide range of distances, and the integration of diverse species, depth ranges, and survey methodologies into a single hierarchical geostatistical model. Studies lacking these features may fail to detect MPA benefits that are otherwise real. The framework we present here unifies inferences on in situ restoration and spillover, offering a template for evaluating spatial protections where responses may be gradual and unevenly distributed among species.

## Acknowledgements

Our heartfelt thanks extend to the many stewardship and technical staff from the Kitasoo Xai’xais, Haíłzaqv, Nuxalk, and Wuikinuxv First Nations who co-lead or contributed to this project (Text S4). The analyses we present here are part of the scientific program of the Central Coast Indigenous Resource Alliance and were funded by the Fisheries and Oceans Canada Ecosystem and Ocean Science Contribution Framework (SFSF-2023-094) and the NSERC Discovery Grant (RGPIN-2025-04787). We thank M.R. Siegle, four anonymous reviewers, and the handling editor for helpful comments on earlier versions of this manuscript.

## Statement of authorship

KW and SC led the analyses, AF led the field collection of data. KW, AF, and SC overviewed model results and interpretations. AF and KW wrote the first draft of the manuscript, and all authors contributed substantially to revisions.

## Data and code availability statement

Data collected for this study are owned and coordinated by the Wuikinuxv, Nuxalk, Kitasoo Xai’xais and Haíłzaqv First Nations, who are maintaining its confidentiality under established principles for Indigenous data governance and sovereignty (First Nations Information Governance Centre 2025). Data may potentially be made available upon request to the corresponding author or to the Central Coast Indigenous Resource Alliance (https://www.ccira.ca/), and an exception to the data sharing policy of the journal was provided by the Editor-in-Chief, Dr. Peter H. Thrall. All model code is archived at https://doi.org/10.5281/zenodo.22166042.

## Notes

https://doi.org/10.5281/zenodo.22166042

### Competing Interest Statement

The authors have declared no competing interest.

### Summary of Updates

Minor revisions to the manuscript after round 2 of peer-review in journal submission.

