## Supplementary material for "Groundfish with diverse life histories increase in size and abundance with proximity to spatial protections": Table S1

**Text S1: History of rockfish over-exploitation and improved management measures in British Columbia.**

Beginning in the early 1980s, many rockfishes in British Columbia (BC), Canada, underwent steep biomass declines that paralleled a period of overexploitation (Yamanaka & Logan 2010). Coinciding with improved fishery management, most declines tapered off in the early 2000s, and biomass for most assessed rockfishes stabilized at levels that can support maximum sustainable yield harvests (Anderson *et al.* 2021). For yelloweye rockfish (*S. ruberrimus*)—a culturally significant species and upper level predator—median body length in the catches of Indigenous fishers declined from 84 cm in the 1980s to 46 cm in the 2010s (Eckert *et al.* 2018). Additionally, fishery independent surveys showed that the mean body sizes of yelloweye rockfish and quillback rockfish (*S. maliger*: another culturally significant species) and the mean age of yelloweye rockfish were declining between the early 2000s and mid 2010s (McGreer & Frid 2017).

To improve the management of rockfish, between 2004 and 2007 Fisheries and Oceans Canada (DFO) implemented 162 Rockfish Conservation Areas, or RCAs (Yamanaka & Logan 2010) throughout BC. RCAs were part of a broader conservation strategy that included accounting for all catch, reductions in fishing mortality, and improved monitoring and stock assessment (Yamanaka & Logan 2010). RCAs are long-term spatial closures that exclude bottom trawl, groundlines, and hook-and-line—the gear types responsible for most fishing mortality of rockfish. RCAs, however, allow mid-water trawls, which catch benthopelagic rockfishes, and invertebrate traps, which catch demersal rockfishes incidentally (Thornborough *et al.* 2020). The locations of RCAs were chosen so that they would cumulatively encompass 30% of good rockfish habitat in the Strait of Georgia and adjacent areas of southern Vancouver Island, and 20% of good rockfish habitats in the remainder of the coast (i.e., allowing 70%-80% of rockfish habitat to remain open to fisheries. Locations containing good rockfish habitat were determined by spatial analyses and stakeholder consultation (Yamanaka & Logan 2010).

RCAs were established at the exclusion of Indigenous governance and knowledge (DFO 2026). Some RCAs, however, are now included in the design of an MPA network being implemented for Canada’s Northern Shelf Bioregion, for which 17 First Nations are co-governance and technical partners with federal and provincial governments (Beaty *et al.* 2024).

Earlier studies in BC failed to produce strong evidence for increased abundances or body sizes of rockfish inside RCAs (Haggarty *et al.* 2016b; McGreer *et al.* 2020). The reasons potentially include the young age of RCAs at the time of data collection (Haggarty *et al.* 2016b), lack of fisher compliance (Haggarty *et al.* 2016a; Iacarella *et al.* 2023), and limitations of the available data and analytical methods (McGreer *et al.* 2020).

**Table S1.** Survey methods used for data collection.

| **Survey method** | **Sampling years** | **Depth, m (mean)** | **Key characteristics** | **Data used in current analyses.** | **Notes** |
| --- | --- | --- | --- | --- | --- |
| Shallow diver transects (Frid *et al.* 2018; McGreer *et al.* 2020) | 2013, 2015–2023 | 5-35 (21) | Belt transects (30 m $\times$ 4 m $\times$ 4 m, or 480 m^3^), along depth contours. | 1. Relative density (count/480 m^3^) of fish and structural corals, by species.  2. Total length of individual fish (see Text S2).  Estimates of habitat structural complexity. | Larger, older rockfishes tend to be deeper than the max. depth of dive surveys. Analyses excluded fish with lengths <20% of *L*_∞_ (see text). |
| Mid-depth video transects (Frid *et al.* 2018, 2019, 2020) | 2015-2018 | 15-200  (67) | Video transects of variable size were divided into bins covering 75-130 m^2^ (mean = 116 m^2^) to reduce depth and habitat variability within spatial units. (Bins <75 m^2^ are end cuts and bins >130 m^2^ reflect GPS data gaps; analyses exclude both.) Parallel laser beams (10-cm apart) provide a distance scale. | 1. Relative density (count/m^2^) of fish.  2. Estimates of habitat structural complexity. | Fish counts were corrected for species detection biases (i.e., attraction to laser beams) (Frid *et al.* 2019, 2020). Camera lacks panning/tilting ability and depth capacity of BOOTS camera (see below). The lower bound for bin size in earlier analyses^2^ was 100 m^2^, which we lowered to 75 m^2^ to not exclude some coral-rich areas. |
| Deep video transects (BOOTS) (Gale *et al.* 2017) | 2018 | 100-500  (253) | Belt transects varied widely in area but were divided into similar size bins, as described for mid-depth video transects. Parallel laser beams (10-cm apart) provide a distance scale. | 1. Relative density (count/m^2^) of fish.  2. Estimates of habitat structural complexity. | Fish counts were corrected for species detection biases (Frid *et al.* 2019, 2020). Transect bins averaged 120 m^2^. |
| Hook-and-line (Frid *et al.* 2016) | 2006-2007; 2013-2015 | 15-205  (57) | Standardized gear fished the bottom for 15-min or 30-min sampling sessions. | 1. Relative density (count/min) for each fish species.  2. Total length of individual fish | During 2006-2007 data were collected by the Heiltsuk Nation prior to CCIRA’s inception. Habitat structural complexity was estimated via a substrate model (Gregr *et al.* 2021). |
| Sampling the landings of Indigenous food fisheries (Frid *et al.* 2016) | 2013-2016 | 15-120  (49) | Researchers either participated in the fishery to collect data, or fishers shared their catch information and lent their “specimens”. | Total length of individual fish | Fishers used baited hooks on longlines (bait type, number of hooks/set and soak times were recorded) or hook-and-line gear with un-baited hooks and a diversity of jigging lures. Habitat structural complexity was estimated via a substrate model (Gregr *et al.* 2021). |

**Text S2**. ***Testing diver accuracy in visual estimates of fish total length.***

Early in their training, divers tested their accuracy measuring fish-shaped models attached to a line suspended in the water column; comparisons of estimated and actual sizes showed low measurement errors that did not vary with model size, and relatively low variation between divers (McGreer *et al.* 2020); this test was applied to divers who estimated almost all fish lengths during 2013-2019. In 2021, a more rigorous test was conducted during 5 surveys in which individual rockfish lengths estimated by divers were compared to the greater accuracy and precision of paired measurements obtained with a stereo camera system (e.g., Stamoulis *et al.* 2020). Stereo camera calibration in the field and post-field image processing used procedures and software described by [SeaGIS](https://www.seagis.com.au/). Correspondence between paired visual and stereo camera measurements was reasonably good (Figs. S1a-c). Neither of the two divers tested, who recorded all fish lengths during 2020-2023, tended to over-estimate fish sizes, but one diver tended to under-estimate fish sizes more than the other (Fig. S1d).

**Figure S1**. Correspondence between paired visual and stereo camera measurements. In panel **d**, “Proportional deviation by diver estimate” is the difference between diver and stereo camera estimates divided by the stereo camera estimate.

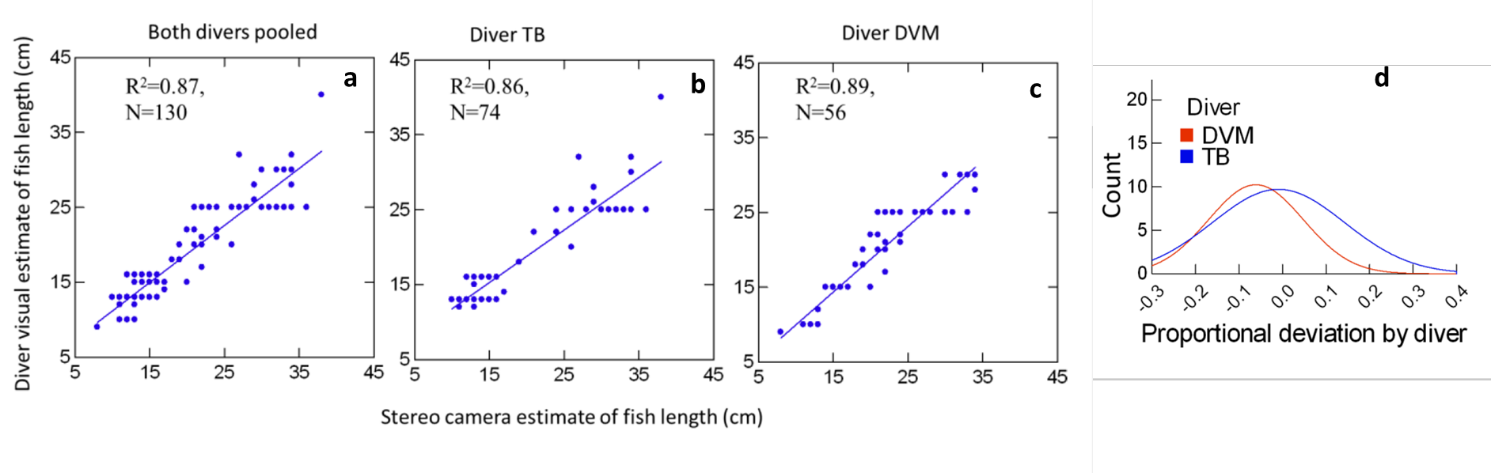

**Text S3. *Causal inference and links between DAG and GLMM statistical model.***

We developed a directed acyclic graph (DAG; Figure S2) to explicitly represent our assumed causal structure (Byrnes & Dee 2025; Correia *et al.* 2026) linking biological responses of total length and CPUE (blue nodes) to the policy treatment of spatial protections (green nodes; RCA proximity and RCA age), additional environmental control variables (white nodes), and unmodelled latent effects (gray nodes). This DAG was then used to guide the implementation of our spatial GLMM in *sdmTMB* (Anderson *et al.* 2025) to ensure our spatial GLMM adjusted for appropriate confounders (Table S2).

The two exposures of interest, RCA proximity and RCA age, were modelled as fixed effects (protection_dist_scale and year_protected_scaled), with interactions allowing the proximity effect to depend on RCA age, pre-RCA cumulative harvest (cumul_catch_before_rca_scaled), and species life history (rmax_scale). Cumulative pre-RCA harvest was an adjusted confounder because RCAs were placed non-randomly with respect to historical fishing intensity, which opens a backdoor path from harvest history into both exposures. Hence, we included this term directly as a fixed effect (cumul_catch_before_rca_scaled) and as an interaction with proximity. We treated cumulative post-RCA harvest as a latent post-treatment mediator in the DAG because conditioning on it could block the causal effects of interest.

Several DAG nodes were addressed through the structure of the data and survey design rather than through the GLMM. The “Depth → Sampling Gear” edge was adjusted within the data because each survey gear's depth range is constrained, and the gear-by-depth confounding is broken by adjusting for both as covariates. Furthermore, the offset log(effort / (1 - chase_prob)) standardised gear-specific catchability so that gears with different effort units were placed on a common ‘catch-per-unit-effort’ scale.

The random spatial field (ω, coded as mesh = barrier_mesh and spatial = "on") and species-specific spatial fields (ε, coded as spatiotemporal = "iid" with time = "species") absorbed unmeasured spatially correlated influences to total length and CPUE (e.g., the “latent spatial factor” node in the DAG). The random slopes for protection_dist_scale varied by rca_name and species, which allowed the effect of RCA proximity to vary across the network of protected areas and across species – this partly accounted for latent species effects, like mobility, and allowed for variable effectiveness from individual RCAs. Sampling gear (gear), structural complexity (complexity_scaled), seasonality (s(biweekly_scale), count model only), and depth (s(log_depth, by = depth_cat)) were included as covariates to control their respective impacts to the body size (total length) and fish density (CPUE) outcomes.

**Figure S2**. Directed acyclic graph (Arif & MacNeil 2023; Byrnes & Dee 2025; Cinelli *et al.* 2024; Pearl 2009) showing hypothesized causal relationships among the outcome (blue), exposure variables (green), non-confounding covariates (white circles), and variables not included in the models (gray). Cumulative harvest after RCA establishment was excluded because it is a post-treatment variable; accounting for it would open a backdoor path. Species mobility was not used because of missing data for 10 species in Table 1. The DAG was constructed with tools provided at <https://dagitty.net/> (Textor *et al.* 2017).

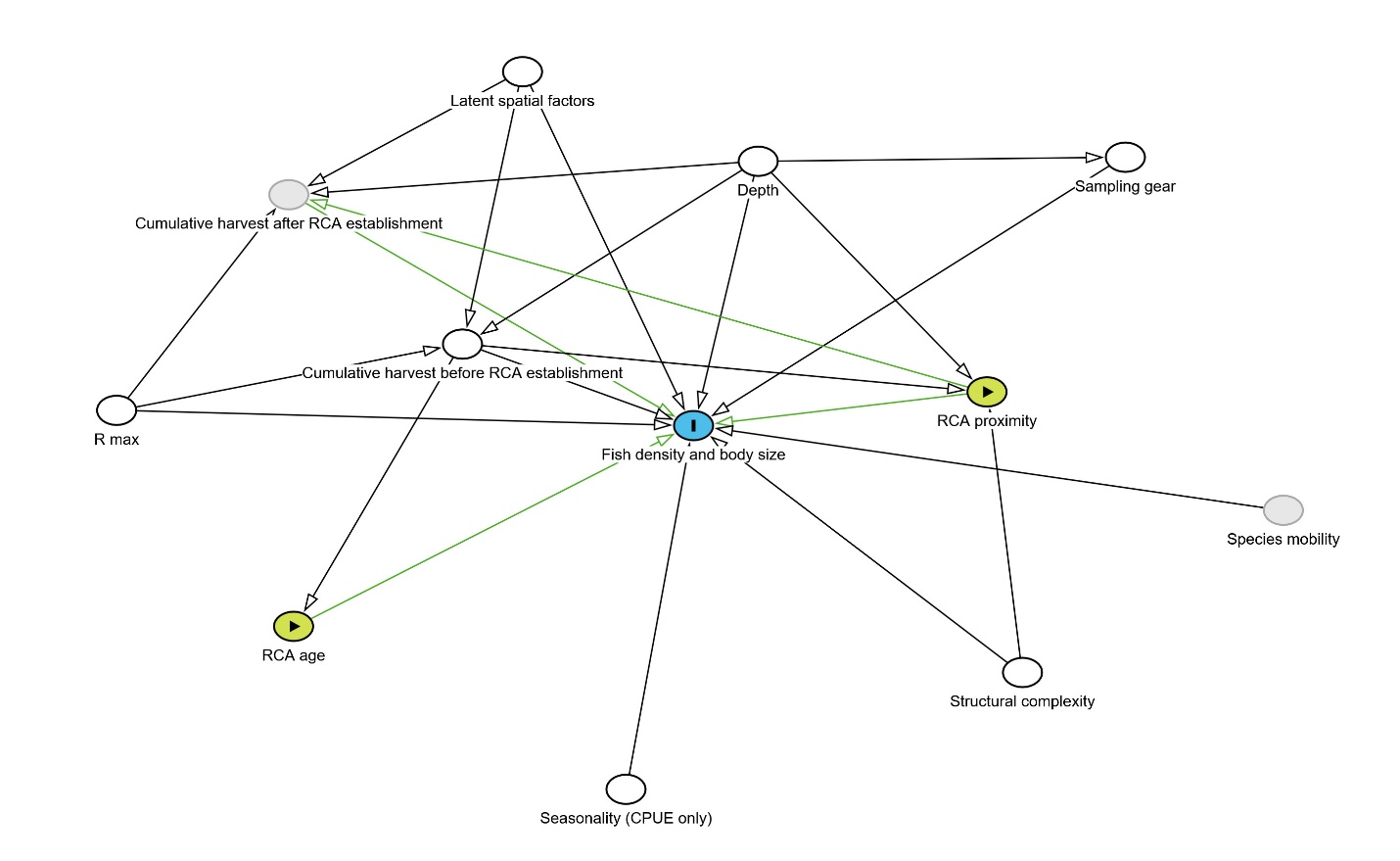

| **Table S2.** Interpreting and mapping each element of the spatial GLMM in *sdmTMB* (Anderson *et al.* 2025) to the directed acyclic graph in Figure S2. With exception of the smoother of log(depth), the model for total length was the same as the CPUE model. Coefficient estimates in Tables S5 and S6. | | |
| --- | --- | --- |
| DAG arrow(s) addressed | GLMM code in *sdmTMB* | Explanation |
| Outcome node: biological response of fish density (CPUE) and body size (total length) | counts ~ | The expected counts of species within a transect is generated by a function of |
| RCA age → Response variables | year_protected_scaled | Effect of RCA age |
| Sampling gear → Fish density and body size | gear | Effect of Survey gear/method |
| Depth → Fish density and body size; Depth → Sampling gear (depth-gear confounding is blocked by adjusting for both) | s(log_depth,by=depth_cat) | Effect of log_e_(depth) (varies by groups of similar species) |
| Structural complexity → Fish density and body size; Structural complexity → RCA proximity (blocked by adjustment) | complexity_scaled | Effect of Habitat complexity |
| Seasonality → Fish density and body size | s(biweekly_scale) | Effect of Seasonality |
| RCA proximity → Fish density and body size | protection_dist_scale | Effect of Proximity to nearest RCA |
| $r_{MAX}$ → Fish density and body size | rmax_scale | Effect of species' life history ($r_{MAX}$) |
| Cumulative harvest before RCA establishment → Fish density and body size; backdoor paths from pre-RCA harvest into RCA age and RCA proximity are blocked by adjustment | cumul_catch_before_rca_scaled | Effect of Cumulative harvest before spatial protection |
| $r_{MAX}$ → Fish density and body size, conditional on RCA age | year_protected_scaled:rmax_scale | Interacting effects of life-history and RCA age |
| RCA proximity → Fish density and body size, conditional on pre-RCA harvest history | protection_dist_scale:cumul_catch_before_rca_scaled | Interacting effects of cumulative harvest and RCA proximity |
| RCA proximity → Fish density and body size, conditional on RCA age | protection_dist_scale:year_protected_scaled | Interacting effects of RCA age on RCA proximity |
| RCA proximity → Fish density and body size, conditional on $r_{MAX}$ | protection_dist_scale:rmax_scale | Interacting effects of life-history on RCA proximity |
| RCA proximity → Fish density and body size (heterogeneous across RCA locations) | (1 + protection_dist_scale \| rca_name) | Intercepts and effects of RCA proximity can vary among RCA locations |
| Species mobility (latent) → Fish density and body size; species-level variation in RCA proximity effect | (1 + protection_dist_scale \| species) | Intercepts and effects of RCA proximity can vary among species |
| Sampling gear → Fish density and body size (effort standardisation) | offset = log(rockfish$effort/(1-rockfish$chase_prob)) | Converts counts into CPUE |
| Sampling gear → Fish density and body size (variance, not mean) | dispformula = ~gear | Variance terms depend on survey gear |
| Latent spatial factors → Fish density and body size (geometric constraint on Matérn covariance) | mesh = barrier_mesh | Latent spatial factors (spatial correlations cannot cross land) |
| Latent spatial factors → Fish density and body size (species-specific) | time = "species" | Latent spatial factors can vary by species |
| Probability distribution of outcome node | family = nbinom2(link="log") | Counts are distributed by negative binomial probability mass function |
| Latent spatial factors → Fish density and body size; Latent spatial factors → Cumulative harvest before/after RCA establishment (absorbs spatially correlated drivers of historical and contemporary fishing pressure) | spatiotemporal = "iid" | Species spatial variation is independent from other species |
| Cumulative harvest after RCA establishment → Fish density and body size (latent mediator) | — | Cumulative harvest post-RCA not included in GLMM because, under our assumed DAG, it blocks causal effects of interest. |
| Depth → Sampling gear | — | Gears have a fixed depth range within a transect. The Depth → Gear pathway does not need adjustment beyond including depth and sampling gear as covariates. |
| Species mobility → Fish density and body size | — | Species mobility differences were not measured directly; the species-level random effects and spatial fields absorb species heterogeneity in responses. |

| **Table S3.** Example of the likelihood weights used in the total-length model for Quillback and Yelloweye rockfish across multiple dive transects that varied in school size. Likelihood weights were each size cohort's share of that species' count in the transect, divided by the mean share within that transect and species. | | | | | | |
| --- | --- | --- | --- | --- | --- | --- |
| Species | Transect | Total length (cm) | No. fish in size cohort | No. fish in transect | Proportion of transect count | Likelihood weight |
| Quillback | A | 10 | 6 | 128 | 0.047 | 0.844 |
| Quillback | A | 10 | 7 | 128 | 0.055 | 0.984 |
| Quillback | A | 10 | 10 | 128 | 0.078 | 1.406 |
| Quillback | A | 12 | 2 | 128 | 0.016 | 0.281 |
| Quillback | A | 12 | 9 | 128 | 0.070 | 1.266 |
| Quillback | A | 14 | 1 | 128 | 0.008 | 0.141 |
| Quillback | A | 15 | 1 | 128 | 0.008 | 0.141 |
| Quillback | A | 15 | 4 | 128 | 0.031 | 0.563 |
| Quillback | A | 15 | 12 | 128 | 0.094 | 1.688 |
| Quillback | A | 15 | 15 | 128 | 0.117 | 2.109 |
| Quillback | A | 15 | 20 | 128 | 0.156 | 2.813 |
| Quillback | A | 18 | 1 | 128 | 0.008 | 0.141 |
| Quillback | A | 20 | 1 | 128 | 0.008 | 0.141 |
| Quillback | A | 20 | 6 | 128 | 0.047 | 0.844 |
| Quillback | A | 22 | 8 | 128 | 0.063 | 1.125 |
| Quillback | A | 22 | 15 | 128 | 0.117 | 2.109 |
| Quillback | A | 25 | 5 | 128 | 0.039 | 0.703 |
| Quillback | A | 25 | 5 | 128 | 0.039 | 0.703 |
| Quillback | B | 20 | 1 | 3 | 0.333 | 1 |
| Quillback | B | 35 | 1 | 3 | 0.333 | 1 |
| Quillback | B | 35 | 1 | 3 | 0.333 | 1 |
| Yelloweye | C | 15 | 1 | 18 | 0.056 | 1 |
| Yelloweye | C | 18 | 1 | 18 | 0.056 | 1 |
| Yelloweye | C | 22 | 1 | 18 | 0.056 | 1 |
| Yelloweye | C | 23 | 1 | 18 | 0.056 | 1 |
| Yelloweye | C | 25 | 1 | 18 | 0.056 | 1 |
| Yelloweye | C | 25 | 1 | 18 | 0.056 | 1 |
| Yelloweye | C | 26 | 1 | 18 | 0.056 | 1 |
| Yelloweye | C | 27 | 1 | 18 | 0.056 | 1 |
| Yelloweye | C | 28 | 1 | 18 | 0.056 | 1 |
| Yelloweye | C | 28 | 1 | 18 | 0.056 | 1 |
| Yelloweye | C | 31 | 1 | 18 | 0.056 | 1 |
| Yelloweye | C | 34 | 1 | 18 | 0.056 | 1 |
| Yelloweye | C | 35 | 1 | 18 | 0.056 | 1 |
| Yelloweye | C | 36 | 1 | 18 | 0.056 | 1 |
| Yelloweye | C | 38 | 1 | 18 | 0.056 | 1 |
| Yelloweye | C | 38 | 1 | 18 | 0.056 | 1 |
| Yelloweye | C | 40 | 1 | 18 | 0.056 | 1 |
| Yelloweye | C | 42 | 1 | 18 | 0.056 | 1 |
| Yelloweye | D | 15 | 1 | 3 | 0.333 | 1 |
| Yelloweye | D | 17 | 1 | 3 | 0.333 | 1 |
| Yelloweye | D | 41 | 1 | 3 | 0.333 | 1 |

| **Table S4.** Totals per transect from examples in Table S3. Likelihood weights sum to the number of size cohorts recorded, so transects recording thousands of fish carry the same total weight as those recording a handful, provided both measured the same number of size classes. | | | | | | |
| --- | --- | --- | --- | --- | --- | --- |
| Species | Transect | Range in total lengths (cm) | No. of size cohorts | No. of fish counted | Sum of weights | Mean weight |
| Quillback | A | 10 – 25 | 18 | 128 | 18 | 1 |
| Quillback | B | 20 – 35 | 3 | 3 | 3 | 1 |
| Yelloweye | C | 15 – 42 | 18 | 18 | 18 | 1 |
| Yelloweye | D | 15 – 41 | 3 | 3 | 3 | 1 |

**Figure S3**. Relationship between relative abundance (ln(CPUE)) and depth among species groups of similar depth preferences.
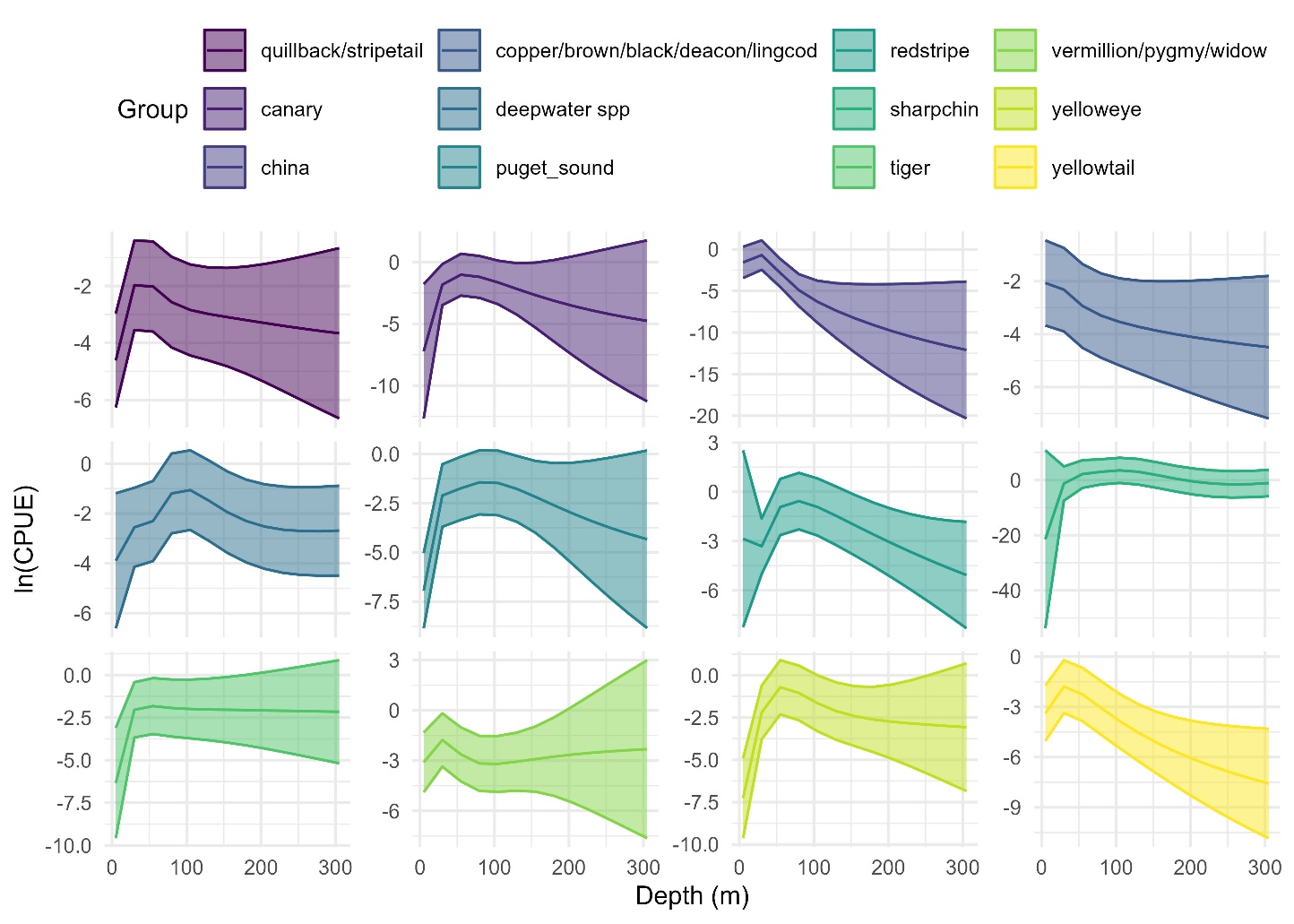

**Figure S4:** Relationship between body size (ln(total length, cm)) and depth among species groups of similar depth preferences.

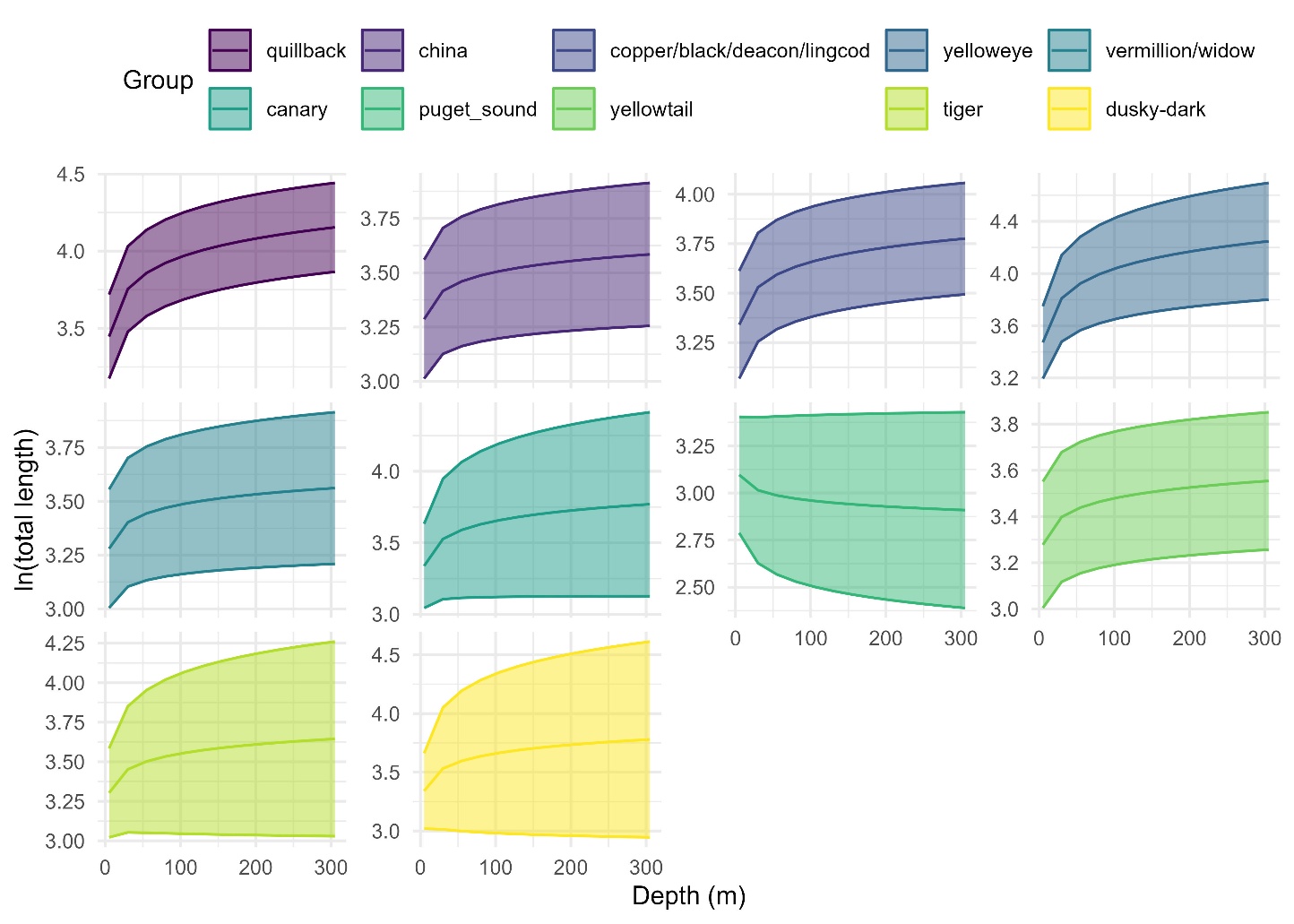

**Figure S5.** Estimates of the main effect of RCA proximity (mean and 95% confidence intervals) under alternative model assumptions of the (a) count model and (b) total length model. Sensitivity tests for the count model included: grouping the dispersion parameter by survey gear or not, including a barrier or non-barrier mesh, testing the NB2 (quadratic variance) versus NB1 (linear variance) parameterization of the negative binomial distribution, resolution of the mesh (cutoff of 2, 3, and 5), and the influence of extreme schooling observations (retained, capped at 3,000 or 1,000 individuals, or excluded). Sensitivity tests for the total length model included: grouping the dispersion parameter by survey gear or not, barrier or non-barrier mesh, Gamma versus lognormal distribution, mesh resolution, and alternative data weighting approaches (base model – weighted by the proportion of fish of that species observed in that size cohort compared to the total number of that species observed in that transect; equal weights per size cohort; weights proportional to expected CPUE, normalised within species; and equal weights per school – where each transect-species group contributes the same total weight regardless of how many fish or size cohorts it contained).
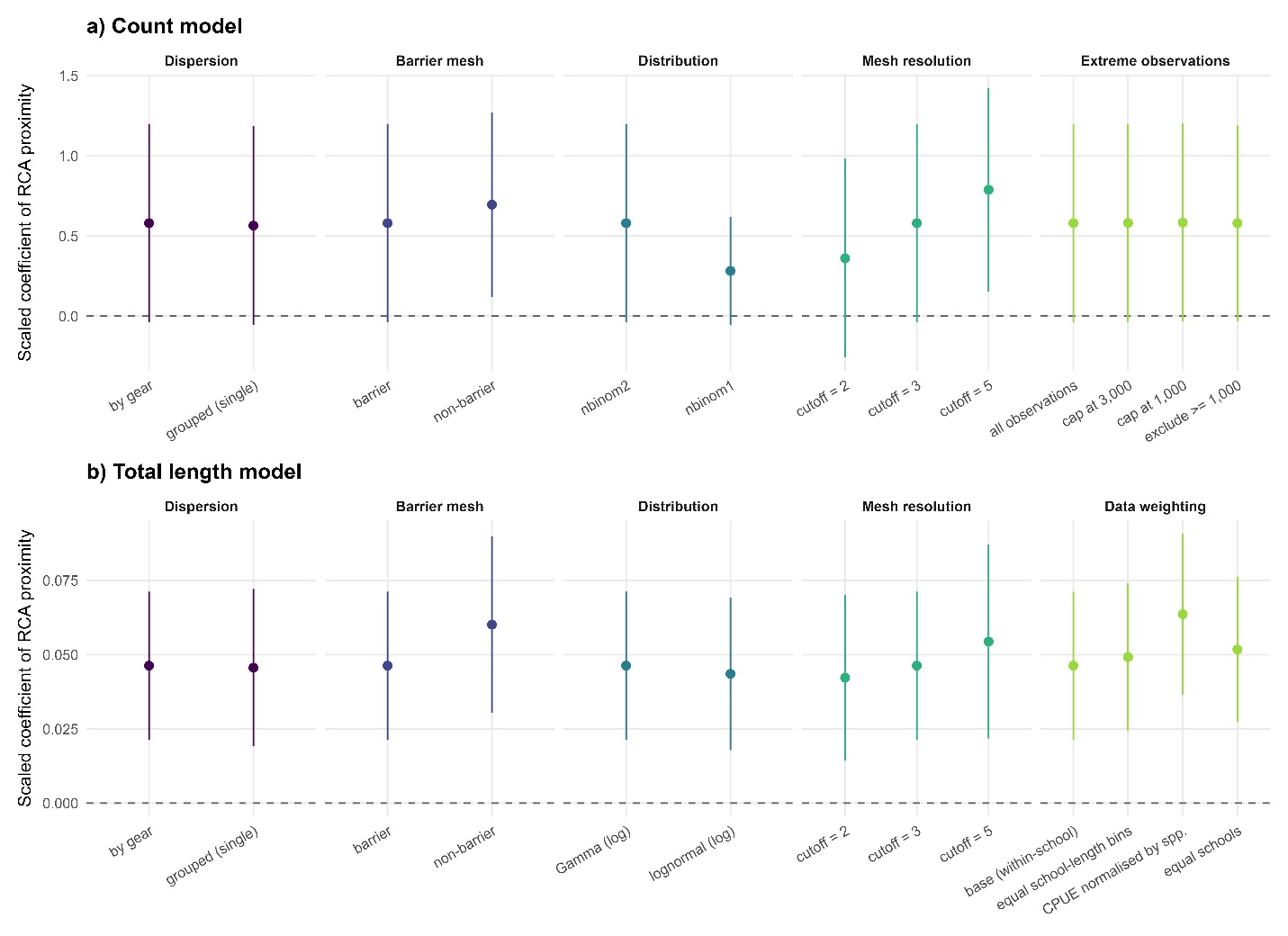

**Figure S6.** Observed versus expected (a) CPUE and (b) total length. Each point is one species-transect record plotted against its own fitted value; dashed lines are 1:1. For CPUE, a constant of 0.001 was added to both axes before log transformation so that zero-catch records remain visible; these form the horizontal band along the lower edge of the panel.

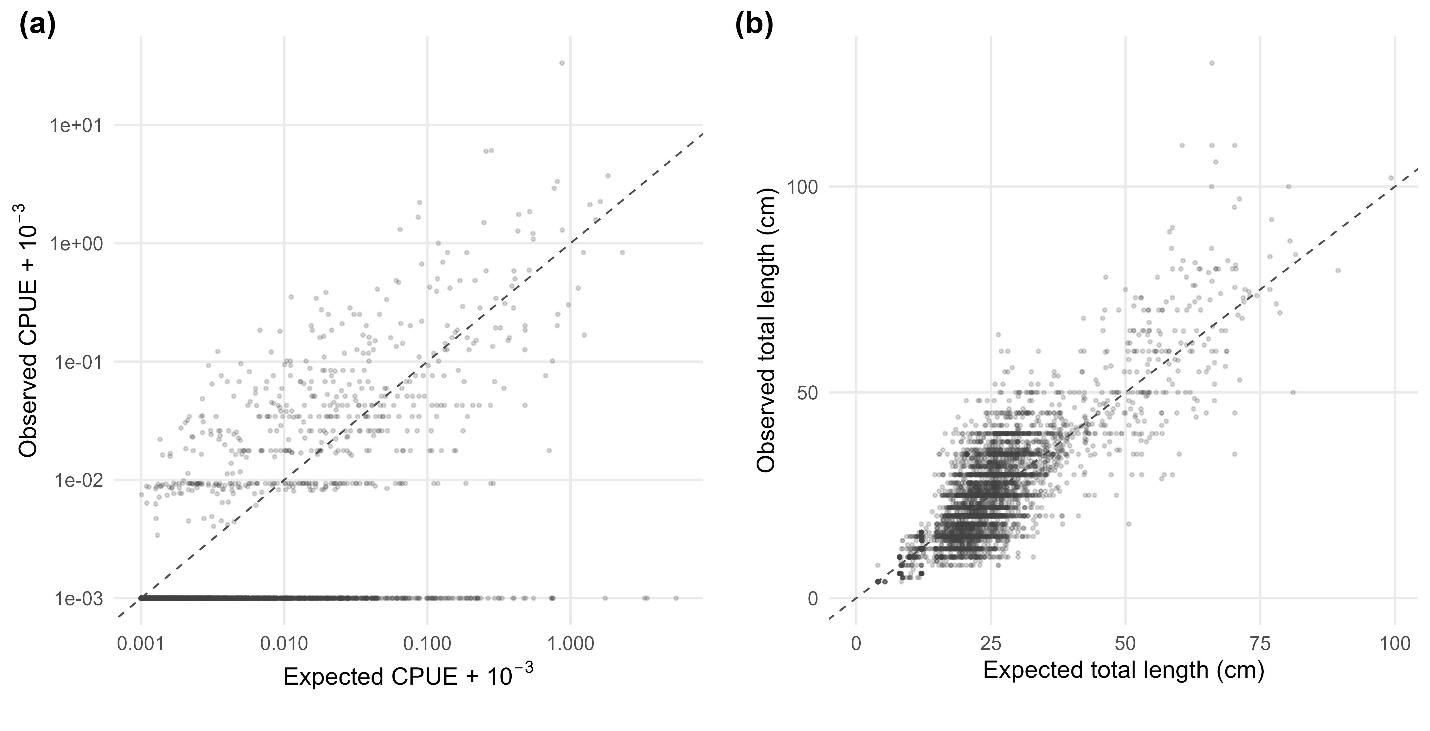

**Figure S7**. Quantile residual diagnostics for spatial GLMMs examining CPUE responses grouped by survey gear. Residuals based on 1,000 simulation draws of observation error with fixed effects held at their maximum likelihood estimates and random effects drawn from the precision matrix of spatial GLMM (Anderson *et al.* 2025; Hartig 2024).

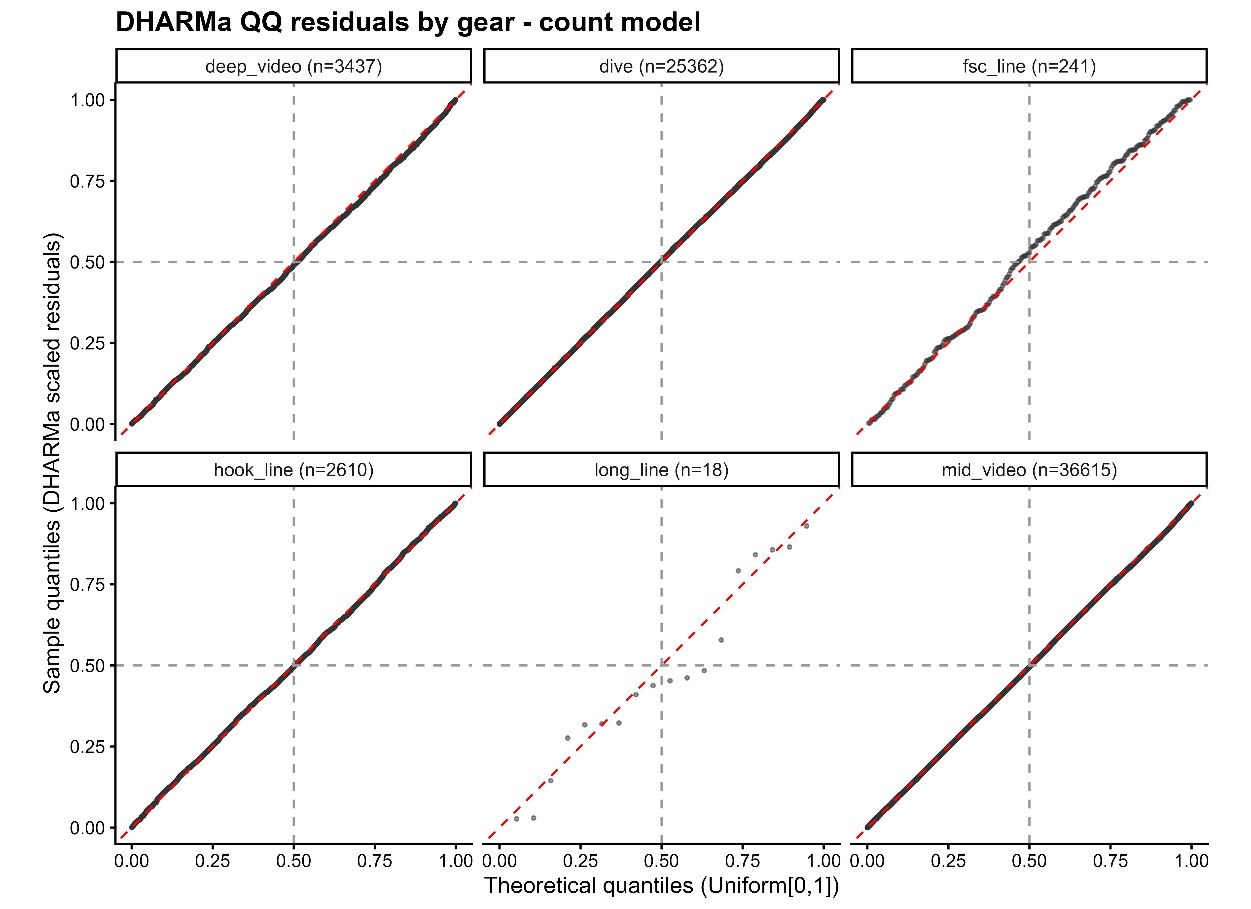

**Figure S8**. Quantile residual diagnostics for spatial GLMMs examining CPUE responses grouped by species. Residuals based on 1,000 simulation draws of observation error with fixed effects held at their maximum likelihood estimates and random effects drawn from the precision matrix of spatial GLMM (Anderson *et al.* 2025; Hartig 2024).

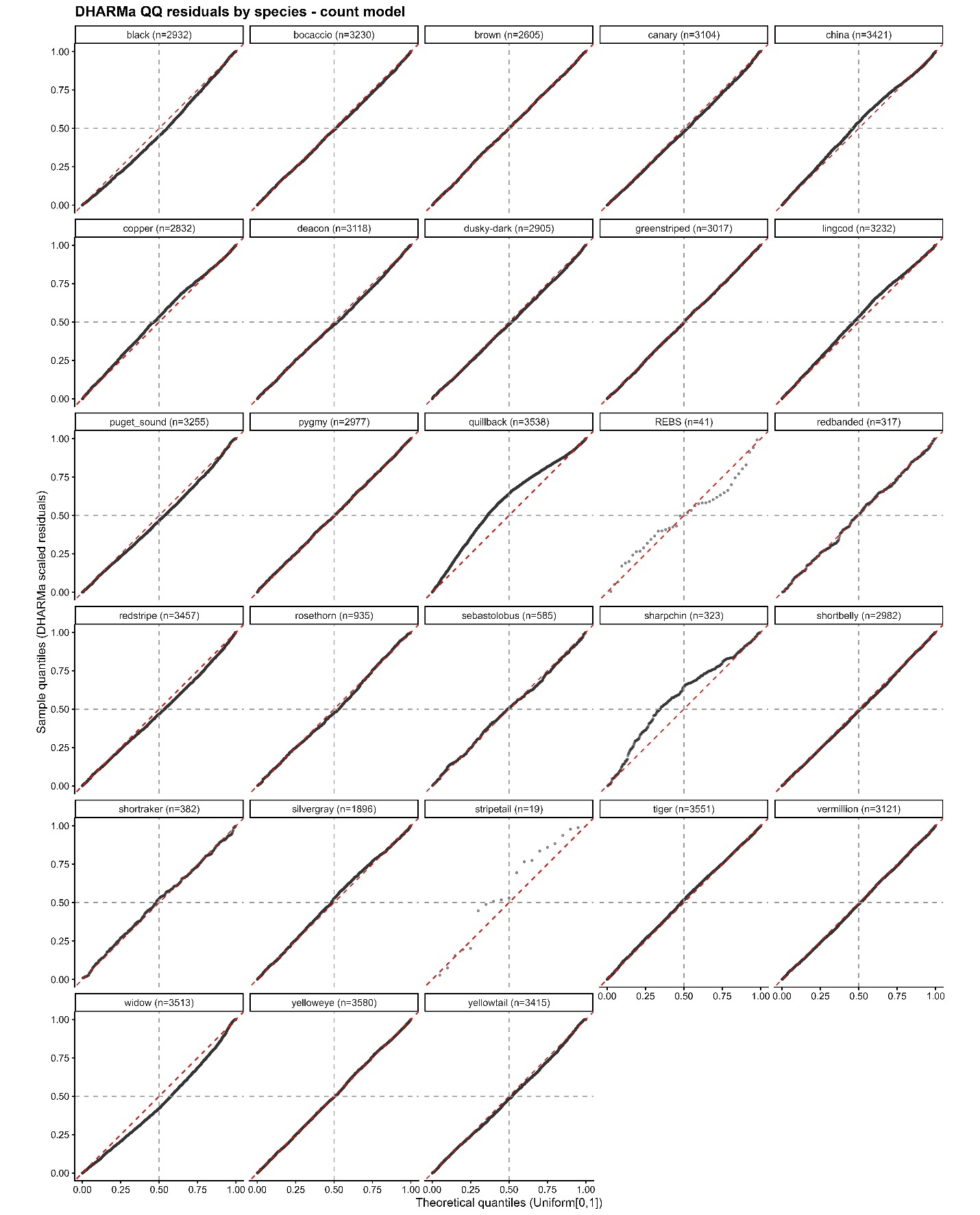

**Figure S9**. Quantile residual diagnostics for spatial GLMMs examining total length responses grouped by survey gear. Residuals based on 1,000 simulation draws of observation error with fixed effects held at their maximum likelihood estimates and random effects drawn from the precision matrix of spatial GLMM (Anderson *et al.* 2025; Hartig 2024).

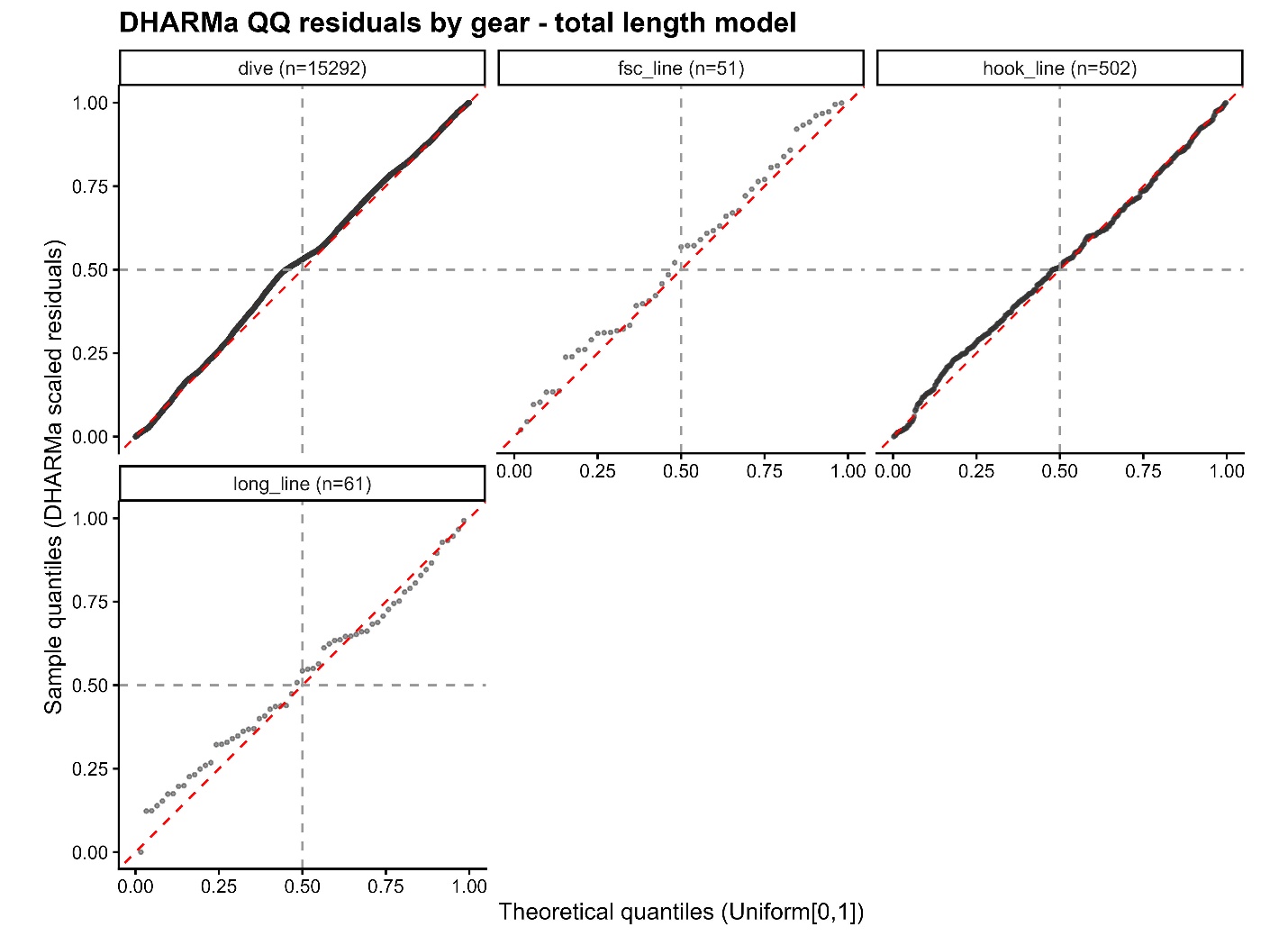

**Figure S10.** Quantile residual diagnostics for spatial GLMMs examining total length responses grouped by species. Residuals based on 1,000 simulation draws of observation error with fixed effects held at their maximum likelihood estimates and random effects drawn from the precision matrix of spatial GLMM (Anderson *et al.* 2025; Hartig 2024).

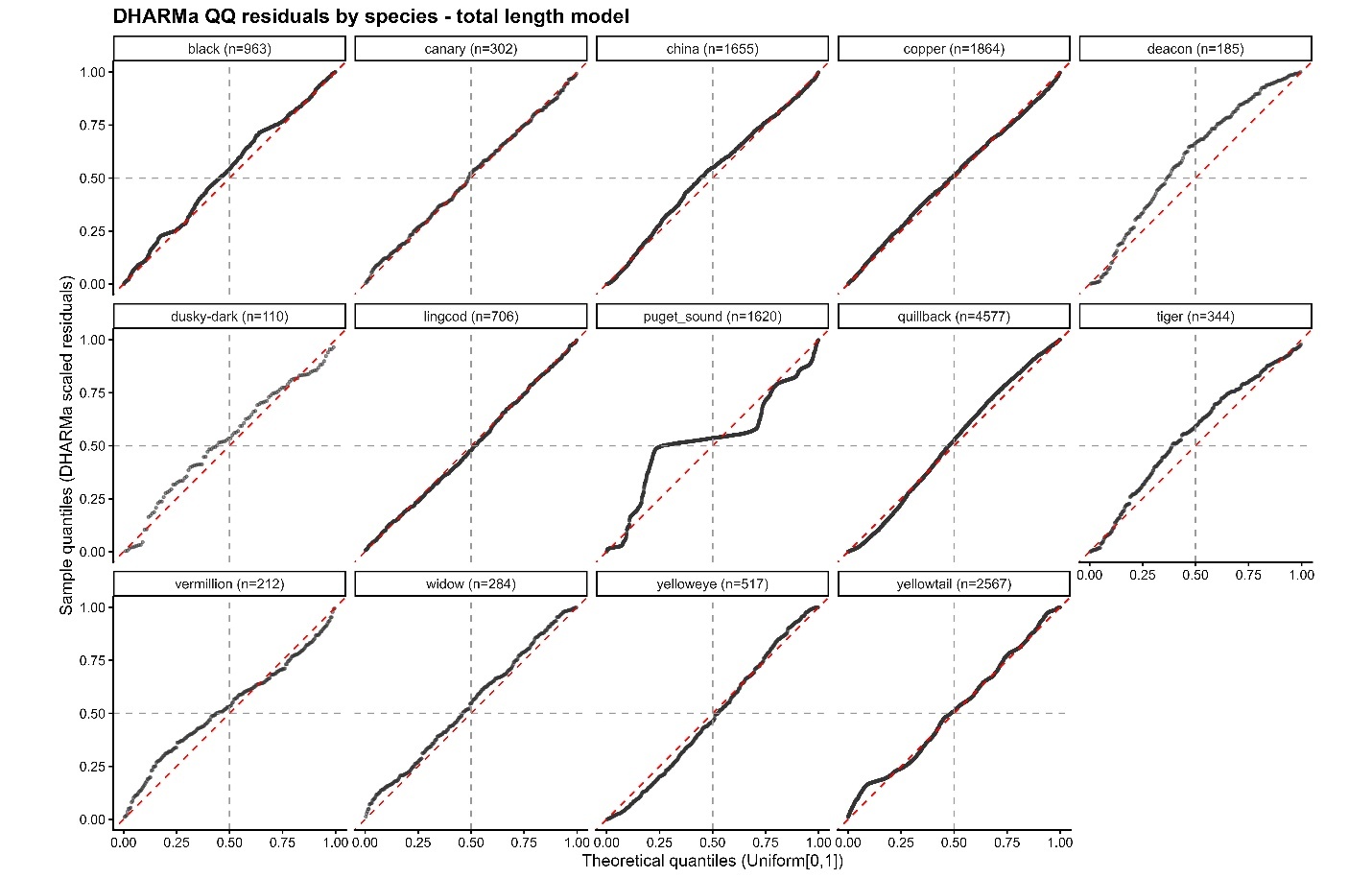

**Figure S11**. Frequency distribution of distances to RCAs at which observations were made.

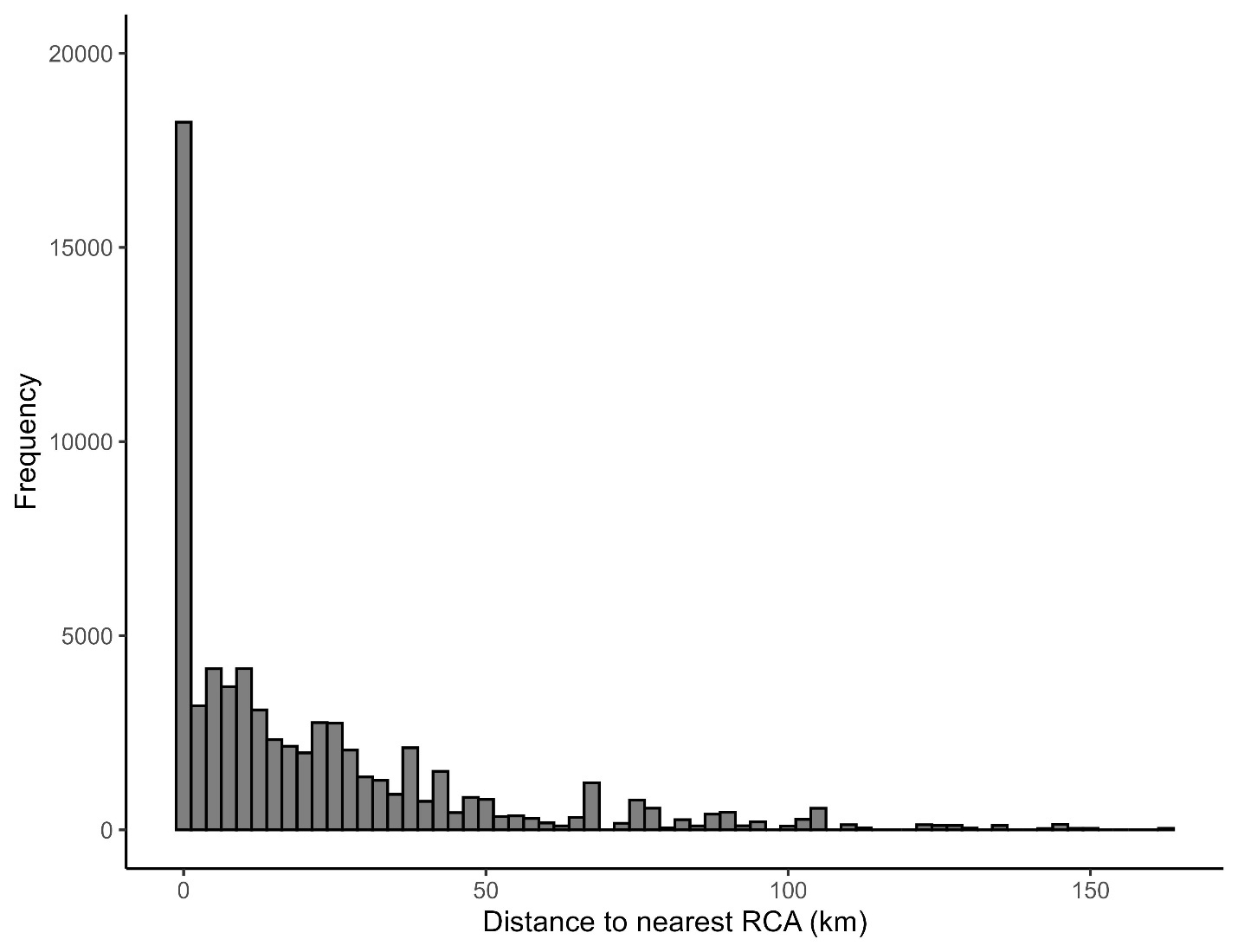

**Figure S12**. Relationship between the von Bertalanffy growth coefficient, *k* (female value), and r_max_ (see Table 1). The solid line and shaded band show the mean and 95% CI, respectively, after excluding Puget Sound rockfish (triangle), which has a substantially higher *k* than any other species; the dashed line shows the fit including all species. Two species with no reported k value (tiger rockfish, shortspine thornyhead) were omitted.

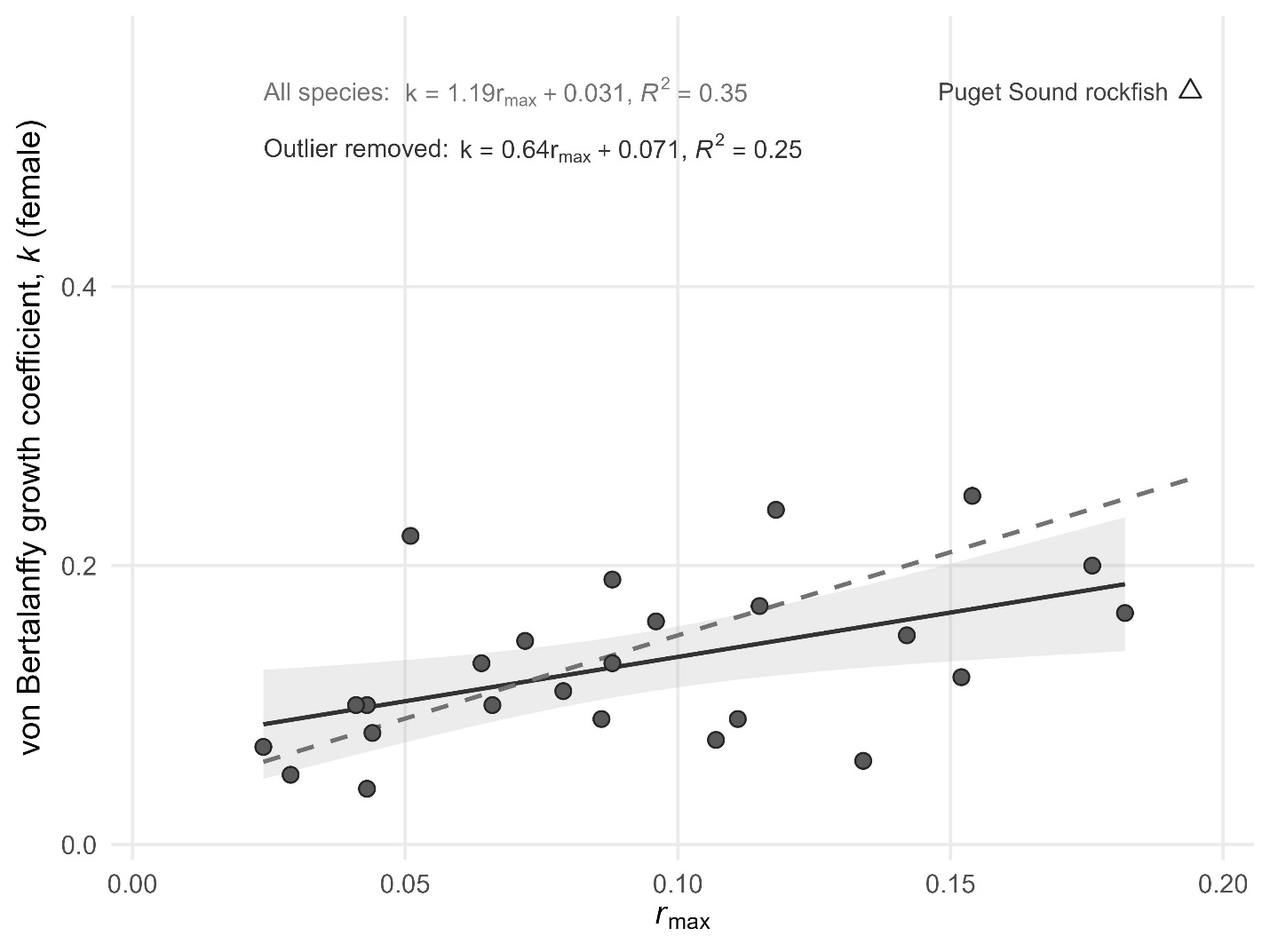

**Figure S13**. Relationships between pre-RCA (1996 to 2004 or 2005) and post-RCA (2004 to year of data collection) commercial fishery harvests by species and category of RCA proximity for CPUE observations. Points represent a survey location with harvest occurring within a 1-km radius. Dotted lines represent 1:1 ratios.

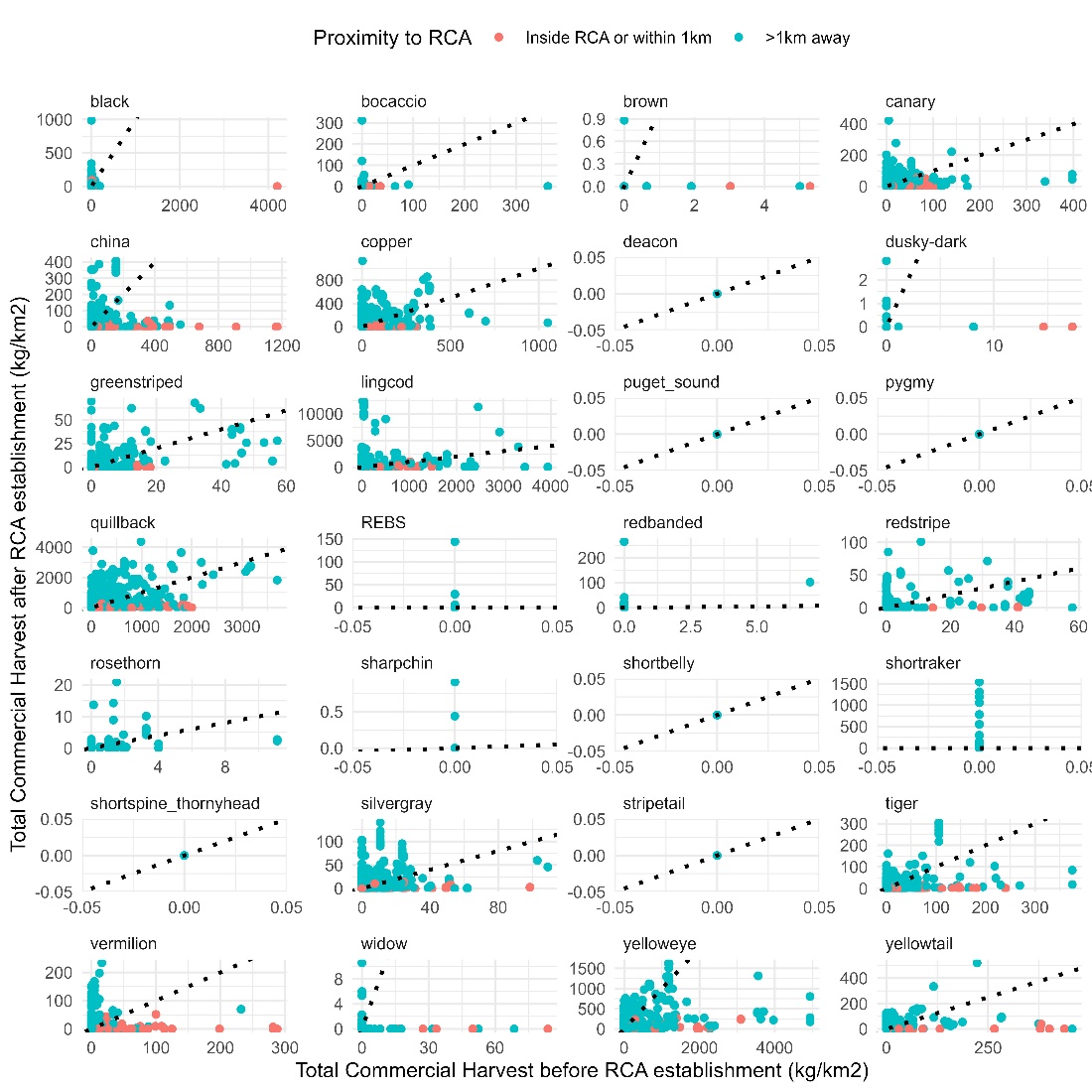

**Figure S14**. Relationships between pre-RCA (1996 to 2004 or 2005) and post-RCA (2004 to year of data collection) commercial fishery harvests by species and category of RCA proximity for body size observations. Points represent a survey location with harvest occurring within a 1-km radius. Dotted lines represent 1:1 ratios.

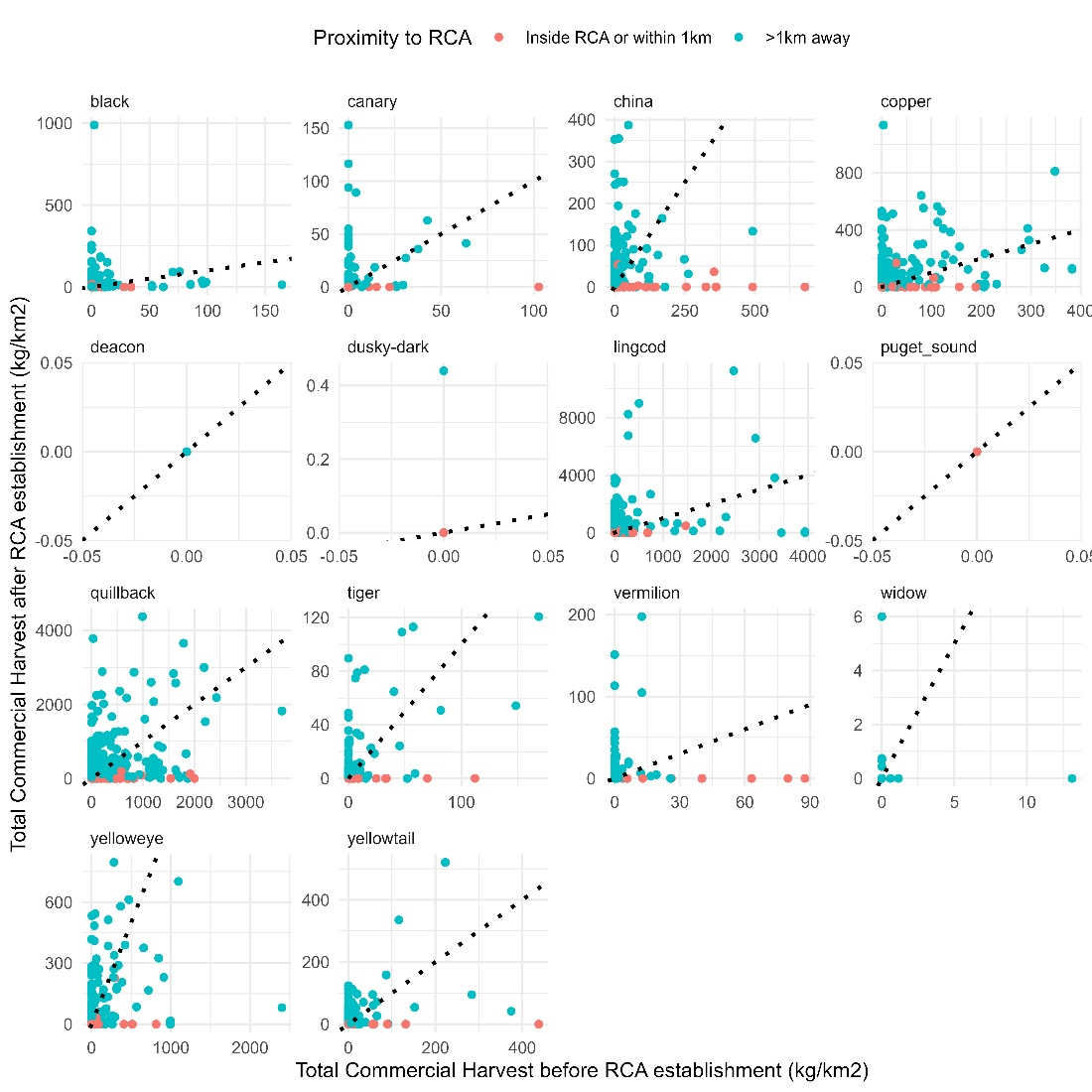

**Figure S15**. Boxplot of RCA distances for each species. The terms REBs represent the rougheye/blackspotted rockfish complex. Dots represent jittered observations spaced horizontally to reflect the density.

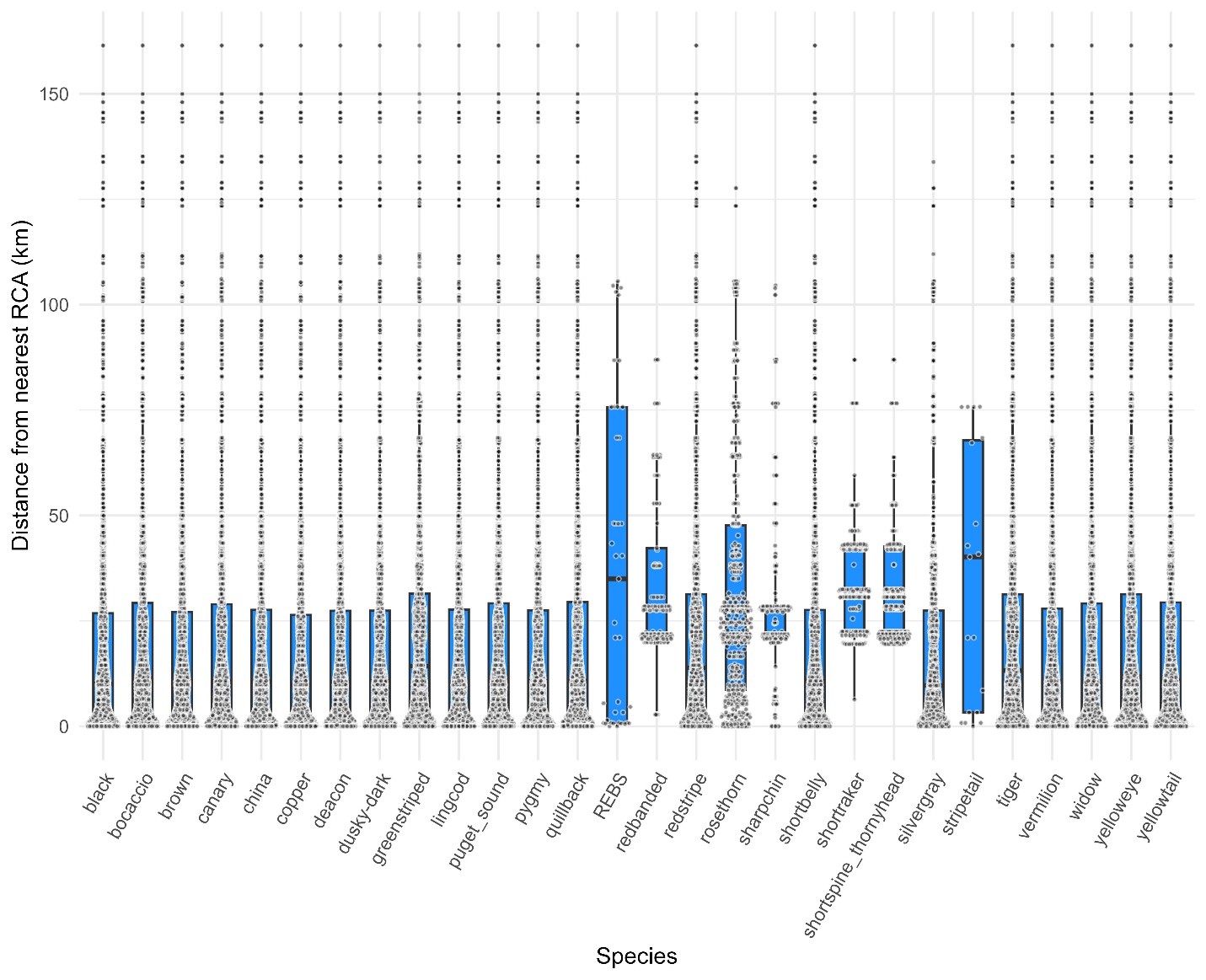

| **Table S5**. Summary of estimated fixed-effect coefficients from spatial generalized linear mixed-effects models of groundfish counts (negative binomial probability distribution with log-link) and total length (Gamma probability distribution with log-link). Pr(β>0) is the proportion of 10,000 simulation draws from the joint precision matrix in which the coefficient exceeded zero, reported as <0.01 or >0.99 where no draws fell on the opposite side. Terms labelled "smoother linear effect" give the linear component of each penalized smoother; the penalized non-linear components are not shown, so these values do not summarise the full depth or seasonal relationships (see Figures S3, S4). | | | | | |
| --- | --- | --- | --- | --- | --- |
| Model | Term | Mean | Std. Err. | 95% CI | Pr(β>0) |
| Count | Intercept | -7.176 | 0.499 | -8.154 – -6.198 | <0.01 |
| Count | annual trend since RCA protection | -0.032 | 0.038 | -0.106 – 0.042 | 0.20 |
| Count | gear effects: deep video | -2.712 | 0.267 | -3.234 – -2.189 | <0.01 |
| Count | gear effects: FSC fishing | 0.116 | 0.418 | -0.702 – 0.935 | 0.61 |
| Count | gear effects: hook-and-line | -1.160 | 0.165 | -1.483 – -0.837 | <0.01 |
| Count | gear effects: long line | -5.183 | 0.555 | -6.272 – -4.094 | <0.01 |
| Count | gear effects: mid-depth video | -2.853 | 0.101 | -3.050 – -2.656 | <0.01 |
| Count | habitat complexity | 0.698 | 0.024 | 0.650 – 0.745 | >0.99 |
| Count | proximity to RCA | 0.581 | 0.316 | -0.038 – 1.199 | 0.97 |
| Count | r_max_ | -0.781 | 0.434 | -1.632 – 0.070 | 0.04 |
| Count | pre-RCA harvest | 0.100 | 0.039 | 0.024 – 0.176 | 0.99 |
| Count | Interaction: annual trend since RCA protection × r_max_ | -0.214 | 0.026 | -0.265 – -0.163 | <0.01 |
| Count | Interaction: RCA proximity × pre-RCA harvest | -0.073 | 0.059 | -0.188 – 0.042 | 0.11 |
| Count | Interaction: RCA proximity × annual trend since RCA protection | 0.063 | 0.037 | -0.010 – 0.136 | 0.95 |
| Count | Interaction: RCA proximity × r_max_ | 0.279 | 0.212 | -0.137 – 0.695 | 0.91 |
| Count | depth smoother linear effect: quillback/stripetail | -0.450 | 1.130 | -2.665 – 1.765 | 0.34 |
| Count | depth smoother linear effect: canary | -0.400 | 3.820 | -7.887 – 7.087 | 0.47 |
| Count | depth smoother linear effect: china | -2.330 | 2.240 | -6.720 – 2.060 | 0.14 |
| Count | depth smoother linear effect: copper/brown/black/deacon/lingcod | -0.610 | 0.630 | -1.845 – 0.625 | 0.17 |
| Count | depth smoother linear effect: deepwater spp. | -17.460 | 6.080 | -29.377 – -5.543 | <0.01 |
| Count | depth smoother linear effect: puget_sound | -2.880 | 2.040 | -6.878 – 1.118 | 0.08 |
| Count | depth smoother linear effect: redstripe | -4.020 | 4.400 | -12.644 – 4.604 | 0.19 |
| Count | depth smoother linear effect: sharpchin | 17.420 | 17.740 | -17.350 – 52.190 | 0.83 |
| Count | depth smoother linear effect: tiger | 0.920 | 2.020 | -3.039 – 4.879 | 0.68 |
| Count | depth smoother linear effect: vermilion/pygmy/widow | -1.980 | 2.660 | -7.194 – 3.234 | 0.22 |
| Count | depth smoother linear effect: yelloweye | -2.060 | 2.620 | -7.195 – 3.075 | 0.22 |
| Count | depth smoother linear effect: yellowtail | -2.130 | 1.090 | -4.266 – 0.006 | 0.02 |
| Count | seasonality smoother linear effect | -0.270 | 0.390 | -1.034 – 0.494 | 0.24 |
| Total length | Intercept | 2.942 | 0.087 | 2.771 – 3.113 | >0.99 |
| Total length | annual trend since RCA protection | -0.020 | 0.006 | -0.032 – -0.008 | <0.01 |
| Total length | gear effects: FSC fishing | 0.205 | 0.041 | 0.124 – 0.285 | >0.99 |
| Total length | gear effects: hook-and-line | 0.178 | 0.018 | 0.143 – 0.214 | >0.99 |
| Total length | gear effects: long line | 0.309 | 0.033 | 0.244 – 0.373 | >0.99 |
| Total length | log(depth) | 0.165 | 0.009 | 0.147 – 0.183 | >0.99 |
| Total length | habitat complexity | 0.014 | 0.003 | 0.008 – 0.020 | >0.99 |
| Total length | proximity to RCA | 0.046 | 0.013 | 0.021 – 0.071 | >0.99 |
| Total length | r_max_ | 0.027 | 0.093 | -0.157 – 0.210 | 0.61 |
| Total length | pre-RCA harvest | 0.010 | 0.004 | 0.002 – 0.019 | 0.99 |
| Total length | log(depth): china | -0.098 | 0.020 | -0.138 – -0.059 | <0.01 |
| Total length | log(depth): copper/brown/black/deacon/lingcod | -0.062 | 0.011 | -0.084 – -0.039 | <0.01 |
| Total length | log(depth): yelloweye | 0.056 | 0.031 | -0.005 – 0.116 | 0.96 |
| Total length | log(depth): vermilion/pygmy/widow | -0.097 | 0.024 | -0.144 – -0.050 | <0.01 |
| Total length | log(depth): canary | -0.041 | 0.055 | -0.148 – 0.066 | 0.23 |
| Total length | log(depth): puget_sound | -0.208 | 0.038 | -0.281 – -0.134 | <0.01 |
| Total length | log(depth): yellowtail | -0.089 | 0.014 | -0.116 – -0.062 | <0.01 |
| Total length | log(depth): tiger | -0.092 | 0.057 | -0.204 – 0.019 | 0.05 |
| Total length | log(depth): dusky-dark | -0.057 | 0.078 | -0.209 – 0.095 | 0.23 |
| Total length | Interaction: annual trend since RCA protection × r_max_ | 0.021 | 0.004 | 0.013 – 0.028 | >0.99 |
| Total length | Interaction: RCA proximity × pre-RCA harvest | 0.000 | 0.006 | -0.012 – 0.011 | 0.48 |
| Total length | Interaction: RCA proximity × annual trend since RCA protection | 0.010 | 0.004 | 0.002 – 0.018 | >0.99 |
| Total length | Interaction: RCA proximity × r_max_ | -0.018 | 0.009 | -0.035 – -0.001 | 0.02 |
| Total length | seasonality smoother linear effect | -0.370 | 0.070 | -0.507 – -0.233 | <0.01 |

| **Table S6.** Summary of estimated variance parameters from spatial generalized linear mixed-effects models of groundfish counts (negative binomial probability distribution with log-link) and total length (Gamma probability distribution with log-link). | | | |
| --- | --- | --- | --- |
| Model | Description | Estimate | 95% CI |
| Count | Matern range (km; distance where spatial correlation ~ 0.13) | 17.192 | 15.463 – 19.114 |
| Count | Spatial random field SD ($\omega$) | 1.214 | 1.011 – 1.457 |
| Count | Spatio-species random field SD ($\epsilon$) | 1.941 | 1.832 – 2.057 |
| Count | Group-level random-intercept SD by RCA name | 0.353 | 0.108 – 1.150 |
| Count | Group-level random-slope SD by RCA name | 0.467 | 0.229 – 0.953 |
| Count | Group-level random-intercept SD by species | 2.218 | 1.663 – 2.960 |
| Count | Group-level random-slope SD by species | 0.989 | 0.694 – 1.409 |
| Count | SD of log(depth) smoother: quillback/stripetail | 6.446 | 3.022 – 13.751 |
| Count | SD of log(depth) smoother: canary | 11.641 | 3.792 – 35.741 |
| Count | SD of log(depth) smoother: china | 10.772 | 4.726 – 24.551 |
| Count | SD of log(depth) smoother: copper/brown/black/deacon/lingcod | 2.977 | 1.411 – 6.282 |
| Count | SD of log(depth) smoother: deepwater spp | 58.116 | 27.463 – 122.981 |
| Count | SD of log(depth) smoother: puget_sound | 10.676 | 4.678 – 24.367 |
| Count | SD of log(depth) smoother: redstripe | 16.257 | 8.167 – 32.363 |
| Count | SD of log(depth) smoother: sharpchin | 66.295 | 20.229 – 217.264 |
| Count | SD of log(depth) smoother: tiger | 7.109 | 2.937 – 17.205 |
| Count | SD of log(depth) smoother: vermilion/pygmy/widow | 16.765 | 9.094 – 30.907 |
| Count | SD of log(depth) smoother: yelloweye | 12.462 | 5.864 – 26.482 |
| Count | SD of log(depth) smoother: yellowtail | 5.645 | 2.903 – 10.978 |
| Count | SD seasonality smoother | 2.143 | 1.050 – 4.373 |
| Count | Negative binomial dispersion parameter ($\theta;$log space) for dive transects | -1.827 | -1.873 – -1.782 |
| Count | Negative binomial dispersion parameter ($\theta;$log space) for deep video | 1.177 | 0.795 – 1.560 |
| Count | Negative binomial dispersion parameter ($\theta;$log space) for FSC fishing | 0.252 | -0.572 – 1.076 |
| Count | Negative binomial dispersion parameter ($\theta;$log space) for hook-and-line | 0.691 | 0.402 – 0.980 |
| Count | Negative binomial dispersion parameter ($\theta;$log space) for long line | 1.343 | 0.221 – 2.465 |
| Count | Negative binomial dispersion parameter ($\theta;$log space) for mid-depth video | -0.277 | -0.372 – -0.182 |
| Total length | Matern range (km; distance where spatial corr ~ 0.13) | 9.755 | 8.126 – 11.709 |
| Total length | Spatial random field SD ($\omega$) | 0.128 | 0.107 – 0.153 |
| Total length | Spatio-species random field SD ($\epsilon$) | 0.135 | 0.125 – 0.147 |
| Total length | Group-level random-intercept SD by RCA name | 0.027 | 0.009 – 0.081 |
| Total length | Group-level random-slope SD by RCA name | 0.013 | 0.004 – 0.045 |
| Total length | Group-level random-intercept SD by species | 0.291 | 0.194 – 0.437 |
| Total length | Group-level random-slope SD by species | 0.018 | 0.009 – 0.039 |
| Total length | SD of seasonality smoother | 0.905 | 0.504 – 1.625 |
| Total length | Observation-error dispersion (Gamma $\phi$log space) for dive transects | 2.463 | 2.440 – 2.486 |
| Total length | Observation-error dispersion (Gamma $\phi$log space) for FSC fishing | 1.044 | 0.523 – 1.566 |
| Total length | Observation-error dispersion (Gamma $\phi$log space) for hook-and-line | 1.377 | 1.228 – 1.526 |
| Total length | Observation-error dispersion (Gamma $\phi$log space) for long line | 2.144 | 1.739 – 2.549 |

| **Table S7.** Summary of the effective degrees of freedom (EDF) for estimated random variables (smoothers, spatial fields, and hierarchical random effects) from spatial generalized linear mixed-effects models of groundfish counts (negative binomial probability distribution with log-link) and total length (Gamma probability distribution with log-link). EDF provides an estimate of the number of equivalent fixed-effect parameters necessary to explain similar amount of variance in the response variable. For random effects, EDF’s closer to 1.0 indicate strong pooling within groups. For smoothers, EDF’s closer to 1.0 indicate the shape is approximately linear. For random fields, EDFs closer to 1.0 indicate smoother surfaces across the domain. | | |
| --- | --- | --- |
| Model | Term | EDF |
| Count | Spatio-species random field SD (epsilon) | 1731.53 |
| Count | Spatial random field SD (omega) | 170.58 |
| Count | Random slopes and intercepts | 53.72 |
| Count | log(depth) smoother: deepwater spp | 7.87 |
| Count | log(depth) smoother: vermillion/pygmy/widow | 6.62 |
| Count | log(depth) smoother: puget_sound | 5.67 |
| Count | log(depth) smoother: quillback/stripetail | 5.55 |
| Count | seasonality smoother | 5.39 |
| Count | log(depth) smoother: yelloweye | 5.11 |
| Count | log(depth) smoother: yellowtail | 4.99 |
| Count | log(depth) smoother: china | 4.85 |
| Count | log(depth) smoother: redstripe | 4.55 |
| Count | log(depth) smoother: sharpchin | 4.22 |
| Count | log(depth) smoother: copper/brown/black/deacon/lingcod | 4.21 |
| Count | log(depth) smoother: canary | 3.33 |
| Count | log(depth) smoother: tiger | 2.91 |
| Count | Total across all groups | 2021.11 |
| Total length | Spatio-species random field SD (epsilon) | 617.28 |
| Total length | Spatial random field SD (omega) | 143.95 |
| Total length | Random slopes and intercepts | 20.05 |
| Total length | seasonality smoother | 7.09 |
| Total length | Total across all groups | 788.36 |

**Table S8.** Species characteristics and number of RCAs with strong responses for count per unit effort (CPUE) and total length (TL). Strong positive and negative responses are operationally defined, respectively, as Pr(β>0) ≥ 0.90 and Pr(β>0) ≤ 0.10 (see Figure 4 in main text). REBS represents rougheye/blackspotted complex.

| Species | No. of RCAs | | | | | L_∞_ | Max harvest (kg/km^2^) | |
| --- | --- | --- | --- | --- | --- | --- | --- | --- |
|  | With observations  CPUE, TL | Positive  CPUE | Negative CPUE | Positive TL | Negative TL |  | *Pre-RCA* | *Post-RCA* |
| Pygmy rockfish | 7, 0 | 0 | 1 | - | - | 21.63 | 0 | 0 |
| Redstripe rockfish | 7, 0 | 0 | 4 | - | - | 37.83 | 58 | 101 |
| Rosethorn rockfish | 7, 0 | 0 | 1 | - | - | 28.76 | 11 | 21 |
| REBS | 4, 0 | 0 | 0 | - | - | 53.04 | 0 | 144 |
| Shortbelly rockfish | 7, 0 | 0 | 1 | - | - | 28.16 | 0 | 0 |
| Greenstriped rockfish | 7, 0 | 0 | 1 | - | - | 34.93 | 57 | 70 |
| Puget Sound rockfish | 7, 7 | 0 | 0 | 0 | 0 | 17.16 | 0 | 0 |
| Silvergray rockfish | 7, 0 | 0 | 1 | - | - | 59.23 | 109 | 140 |
| Stripetail rockfish | 5, 0 | 0 | 0 | - | - | 33.06 | 0 | 0 |
| Yellowtail rockfish | 7, 7 | 0 | 3 | 6 | 0 | 56.74 | 457 | 521 |
| Redbanded rockfish | 3, 0 | 1 | 0 | - | - | 53.33 | 7 | 265 |
| Shortraker rockfish | 3, 0 | 0 | 0 | - | - | 67 | 0 | 1539 |
| Yelloweye rockfish | 7, 7 | 1 | 0 | 7 | 0 | 65.63 | 4,938 | 1,634 |
| Shortspine thornyhead | 3, 0 | 2 | 0 | - | - | 47.39 | 0 | 0 |
| Bocaccio | 7, 0 | 1 | 0 | - | - | 81.33 | 362 | 313 |
| Copper rockfish | 7, 7 | 2 | 0 | 7 | 0 | 45.84 | 1,052 | 1,135 |
| Dusky/dark complex | 7, 7 | 2 | 0 | 7 | 0 | 47.96 | 17 | 3 |
| Brown rockfish | 7, 0 | 1 | 0 | - | - | 51.46 | 5 | 1 |
| Sharpchin rockfish | 7, 0 | 1 | 0 | - | - | 35.73 | 0 | 1 |
| Widow rockfish | 7, 7 | 3 | 0 | 7 | 0 | 54.83 | 84 | 11 |
| Quillback rockfish | 7, 7 | 5 | 0 | 6 | 0 | 39.83 | 3,699 | 4,370 |
| Black rockfish | 7, 7 | 7 | 0 | 7 | 0 | 49.02 | 4,198 | 988 |
| Canary rockfish | 7, 7 | 7 | 0 | 6 | 0 | 58.63 | 397 | 424 |
| China rockfish | 7, 7 | 5 | 0 | 7 | 0 | 34.77 | 1,169 | 408 |
| Deacon rockfish | 7, 7 | 7 | 0 | 7 | 0 | 37.82 | 0 | 0 |
| Lingcod | 7, 7 | 5 | 0 | 7 | 0 | 114.43 | 3,940 | 12,336 |
| Tiger rockfish | 7, 7 | 7 | 0 | 7 | 0 | 47.40 | 375 | 305 |
| Vermilion rockfish | 7, 7 | 7 | 0 | 7 | 0 | 67.11 | 288 | 234 |

**Text S4**. Extended acknowledgements.

We dedicate this manuscript to the memories of Ernie Mason (Kitasoo Xai’xais Nation, key member of the dive survey team and Hereditary Chief, deceased 2023), Hai’mas “Charlie” Mason (Kitasoo Xai’xais Nation, fisher and Hereditary Chief, deceased 2025) and Valerie Shaw (Wuikinuxv Nation, fisher, deceased 2024) who generously shared their knowledge of groundfish. We thank the Nuxalk, Kitasoo/Xai’xais, Haíłzaqv, and Wuikinuxv First Nations for their leadership during the many years of the project, especially Mike Reid (Haíłzaqv), Doug Neasloss (Kitasoo Xai’xais), Abúk Danielle Shaw (Wuikinuxv), and Megan Moody (Nuxalk). The following is a partial list of people who contributed to field data collection: Ernie Mason, Sandie Hankewich, Tristan Blaine, Derek Van Maanen, Andrew McCurdy, John Sampson, Roger Harris, Ernie Tallio, Chris Corbett, Brian Johnson, Charles Saunders, Alec Willie, Gord Moody, Jordan Wilson, Robert Johnson, Randy Carpenter, Richard Reid, Davie Wilson, Julie Carpenter, Doug Neasloss, Vern Brown, Courtney Edwards, Neha Acharya-Patel, Wayne Jacob, Natalie Ban, Lauren Eckert, and Twyla Frid. Additional field contributors and financial support for CCIRA’s field research are acknowledged in earlier publications (see Table S1). Jennifer Long, Darienne Lancaster, and Ian Murdoch contributed to video annotation. Julie Beaumont estimated distances to RCAs. Andy Lamb and Milton Love advised on species identifications from video captures. Dustin Marshall provided helpful insights on the consequence of hyperallometric fecundity in response to spatial protections.
